# Assembly of higher-order SMN oligomers is essential for metazoan viability and requires an exposed structural motif present in the YG zipper dimer

**DOI:** 10.1101/2020.11.07.336172

**Authors:** Kushol Gupta, Ying Wen, Nisha S. Ninan, Amanda C. Raimer, Robert Sharp, Ashlyn M. Spring, Kathryn L. Sarachan, Meghan C. Johnson, Gregory D. Van Duyne, A. Gregory Matera

## Abstract

Protein oligomerization is one mechanism by which homogenous solutions can separate into distinct liquid phases, enabling assembly of membraneless organelles. Survival Motor Neuron (SMN) is the eponymous component of a large macromolecular complex that chaperones biogenesis of eukaryotic ribonucleoproteins, and localizes to distinct membraneless organelles in both the nucleus and cytoplasm. SMN forms the oligomeric core of this complex, and missense mutations within its YG box domain are known to cause Spinal Muscular Atrophy (SMA). The SMN YG box utilizes a variation of the glycine zipper motif to form dimers, but the mechanism of higher-order oligomerization remains unknown. Here, we use a combination of molecular genetic, bioinformatic, biophysical, and computational approaches to show that formation of higher-order SMN oligomers depends on a set of YG box residues that are not involved in dimerization. Mutation of key residues within this new structural motif restricts assembly of SMN to dimers and causes locomotor dysfunction and viability defects in animal models.

## Introduction

Oligomeric proteins represent a significant fraction of cellular proteomes in all three domains of life. Self-interaction (homo-oligomerization) is a widespread and well-established feature of soluble proteins, occurring within a majority of known macromolecular assemblies [1]. Protein oligomerization provides several functional benefits (reviewed in [2, 3]), not least of which is the potential for forming novel and multivalent interaction surfaces that are not present in the monomer. Oligomerization can thus alter protein stability, enzymatic activity, and allosteric interactions; indeed, the manifold opportunities for adding new layers of regulation are too numerous to list.

Protein oligomerization is thought to underlie a phenomenon that allows homogenous solutions of macromolecules to separate, or ‘demix,’ into two co-existing liquid phases [4]. This biomolecular condensation makes it possible to create cellular sub-organelles (i.e. membraneless domains or compartments) with elevated protein concentrations that serve to accelerate biochemical reactions or to sequester key factors away from the cellular milieu [5, 6]. As with most things in nature, the positive benefits provided by multimerization come with a down-side: oligomeric proteins also have the potential to form dysfunctional or pathogenic aggregates [7–10]. Molecular mechanisms underlying a wide variety of physiological and pathological processes are thus being re-examined through the lens of liquid-liquid phase separation [11].

The Survival Motor Neuron (SMN) protein forms the oligomeric core of a multiprotein complex that chaperones the biogenesis of small nuclear ribonucleoproteins (snRNPs) required for pre-mRNA splicing [12, 13]. Together with its Gemin protein partners [14], the SMN complex is also thought to participate in many other important cellular processes, including RNA transport, translation, endocytosis, cytoskeletal maintenance, and intracellular signaling [15–18]. Notably, SMN is a key component of RNP-rich, phase-separated cellular domains known as stress granules [19] and Cajal bodies [20]. Homozygous mutation or deletion of the human *SMN1* gene causes a devastating neuromuscular disease called Spinal Muscular Atrophy (SMA) [21, 22]. Most SMN orthologs have three conserved domains: an N-terminal region responsible for binding to Gemin2, a centrally-located Tudor domain important for binding to Sm-class splicing factors, and a C-terminal YG box region that mediates self-interaction [23, 24]. There are two paralogous copies of *SMN* in humans, *SMN1* and *SMN2* [21]. Non-human primate and other eukaryotic genomes have only a single-copy *Smn* gene [25], deletion of which is lethal in every species studied to date [26]. Note that the Tudor domain of SMN is not essential for eukaryotic viability, as this domain is missing in certain phyla (e.g. the fungal and trypanosomal SMN proteins). However, the YG box is present in all SMN orthologs identified to date and this domain is also important for the protein to associate with Gemin3 and Gemin8 [27–29].

The extent to which defects in SMN oligomerization contribute to the pathophysiology of SMA is not yet known. Roughly half of all SMA-causing missense mutations in *SMN1* are located within the YG box [30], and the predominant protein isoform expressed from the *SMN2* gene contains a truncation of this domain [31]. Despite the widely held belief that oligomerization of SMN is important for its function, there is no direct evidence that higher-order (n > 2) multimers are actually necessary for SMN to carry out its activities. To better understand how the SMN protein forms oligomers, we carried out a detailed structurefunction analysis of the YG box self-interaction domain. Using a broad spectrum of experimental approaches and model systems, we find that formation of higher-order SMN oligomers depends on specific YG box residues that are not involved in dimerization. These identified residues constitute a second structural motif that is not only required for SMN oligomerization *in vitro*, but also for organismal viability, longevity, and locomotor function *in vivo.*

## Results

### Sequence and structural conservation of the SMN YG box

A phylogenetic comparison of the C-terminal domain of diverse SMN orthologs reveals several highly conserved sequence features. In addition to the overall hydrophobic character of this domain, there are three overlapping motifs (Fig. 1A). The G-motif (GxxxGxxxG) is a signature of certain glycine zipper transmembrane proteins that form coiled-coil dimers and oligomers [32]. The s-motif denotes the presence of small amino acid residues (typically Ser, Ala, Thr). The Y-motif (YxxxYxxxY), on its own, is common among unstructured regions of tyrosine-rich RNA binding proteins. However, the interdigitation of these highly conserved tyrosine and glycine residues was shown to form a unique structural variant of the glycine zipper motif, termed a YG zipper [33].

**Figure 1:**
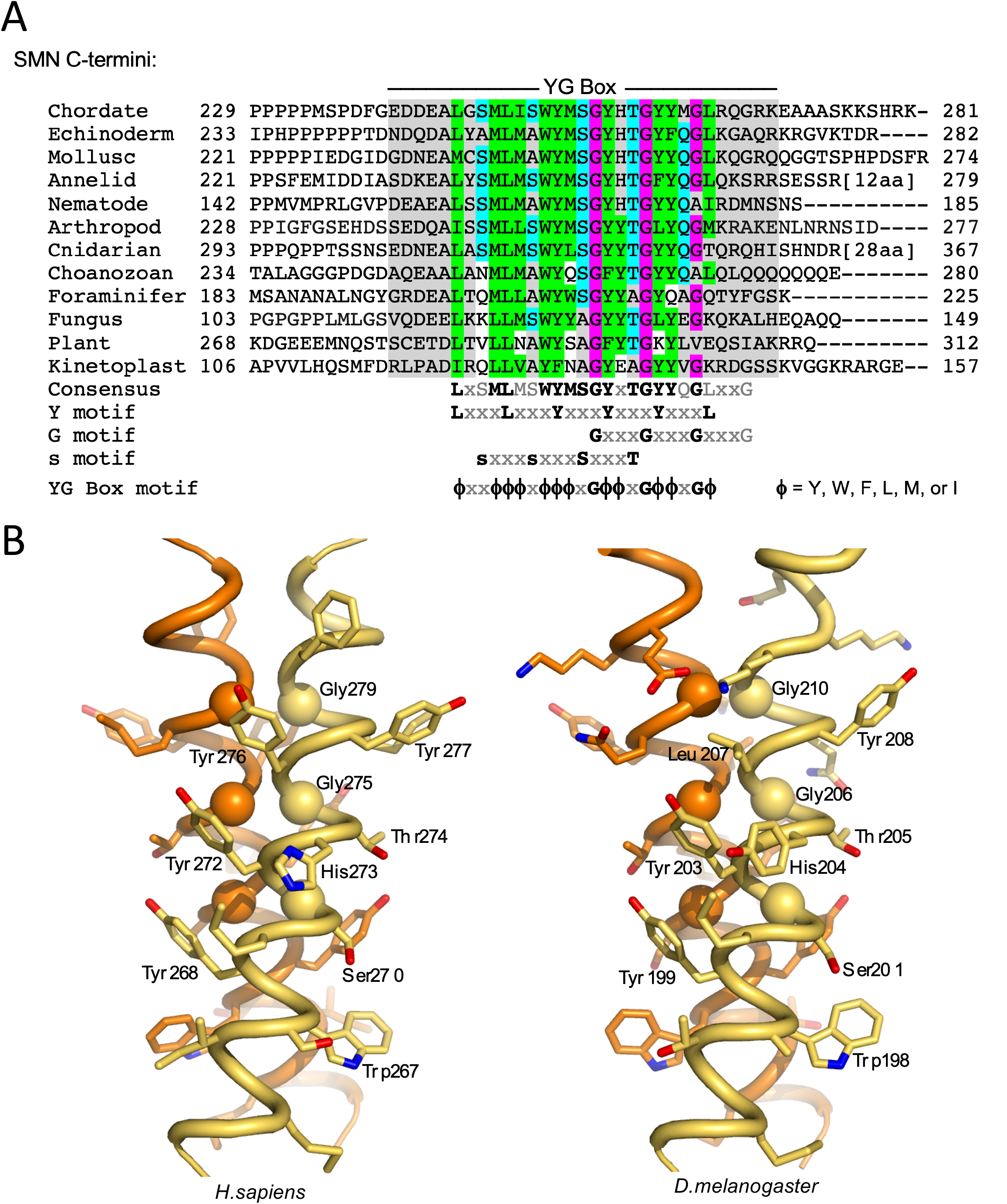
Structural conservation of the SMN YG box domain. ***A.*** Phylogenetic analysis of SMN C-termini from a diverse selection of eukaryotes. Conserved glycine residues are shaded in magenta, hydrophobic residues are in green, and polar residues in teal. Note the regularized spacing of residues in three overlapping motifs (Y, G and s) that are contained within the overall YG box consensus. ***B.*** Structure of the SMN YG-box dimer. Experimental atomic structures of the human (PDB 4GLI) and fission yeast (PDB 4RG5) SMN dimers were used to generate extended models of the human (left) and fruitfly (right) proteins. The three well-conserved glycine residues are shown as Cα spheres. Atomic views were rendered using the program PYMOL [74]. See supplemental Fig. S1 for a more extensive phylogenetic comparison along with additional ultrastructural models.

Within the SMN YG box dimer structure (Fig. 1B), each of the three Y-motif side chains packs against the main chain atoms of the *i+3* glycine residue on the opposing helix. Face-to-face apposition of the three glycine residues with their counterparts in the partner helix leads to a more intimate interface between helix backbones than is found in canonical coiled-coil dimers. This resulting network of intersubunit interactions in SMN is conserved from yeast to human [33, 34]. The broad conservation of SMN YG box sequence features among other eukaryotic groups (e.g. in *Amoebozoa, Plantae, Rhizaria* and *Excavata)* strongly supports the importance of these intersubunit Y-G interactions in dimer formation (Figs. 1A and S1A). As shown in Figs. 1B and S1B, models of the fruitfly and nematode YG zipper dimers can be readily generated from the human and yeast X-ray structures, without steric clashes or need for changes to the main chain helix conformation.

The SMN · Gemin2 heterodimer (SMN·G2) is thought to be the fundamental structural component of the SMN complex, and this subunit is known to form higher-order multimers *in vitro* [33, 34]. Human Gemin2 is monomeric, and does not oligomerize, either alone or in the presence of pre-formed SMN·G2 dimers [34]. For our purposes here, an oligomer of SMN · G2 refers to a species with a stoichiometry of (SMN ·G2)_n_, where n = 2 for a dimer, n = 4 for a tetramer, etc. Notably, purified recombinant SMN [35], as well as those complexes isolated from animal cells [36, 37], are of a size that is much larger than that predicted for a dimer. Yet we currently have no structural models for SMN higher-order oligomerization. Thus, understanding the cellular circumstances and molecular mechanisms whereby SMN assembles into higher-order oligomers is crucial.

### Biophysical properties of metazoan SMN·Gemin2 complexes are conserved

*In vitro* and *in vivo*, human SMN forms complexes of unusually large hydrodynamic size [35]. Size-exclusion chromatography coupled with multi-angle light scattering (SEC-MALS), analytical ultracentrifugation (AUC) and small-angle X-ray scattering (SAXS) data have shown that *Homo sapiens* (hs)SMN ·G2 and hsSMNΔexon5 (hsSMNΔ5) ·G2 complexes exist within a temperature-dependent tetramer to octamer equilibrium with little presence of dimers [34]. For reasons unknown, complexes from fission yeast *Schizosaccharomyces pombe* (spSMN·G2) exist *in vitro* primarily as dimers and tetramers, with no evidence of octamers, or temperature dependence [34]. Furthermore, estimates of SMN protein concentration within living fission yeast cells [38] suggest that the concentration of the dimer (~15 nM) is far below the measured dissociation constant for the tetramer (~1 μM) [34]. Hence, the spSMN·G2 dimer is likely the most abundant species *in vivo* and is presumed to be the basal functional complex required for cell viability [15].

To determine whether the biophysical properties of metazoan SMN proteins are conserved, we generated SMN·G2 complexes from the fruitfly *Drosophila melanogaster* (dmSMN · G2) and nematode *Caenorhabditis elegans* (ceSMN · G2) and carried out a variety of biophysical measurements. As shown in Fig. 2, SEC-MALS shows that fruitfly and nematode SMN·G2 complexes mirror those of the human complex at room temperature. The weight-averaged molecular masses (Mw) of the complexes derived from MALS (Fig. 2A) are consistent with a tetramer-octamer distribution and the sedimentation velocity (SV) profiles obtained via AUC (Fig. 2B) are similar in size and breadth to those previously determined for the hsSMN ·G2 [34]. The sedimentation coefficients of the human complex were also shown to be temperature-dependent, with a shift towards smaller species observed at 4°C. This behavior was recapitulated with the nematode and fly complexes in SV-AUC analysis (Fig. 2B). The temperature dependence of metazoan SMN oligomerization implies that the larger multimers are stabilized primarily by hydrophobic interactions.

**Figure 2:**
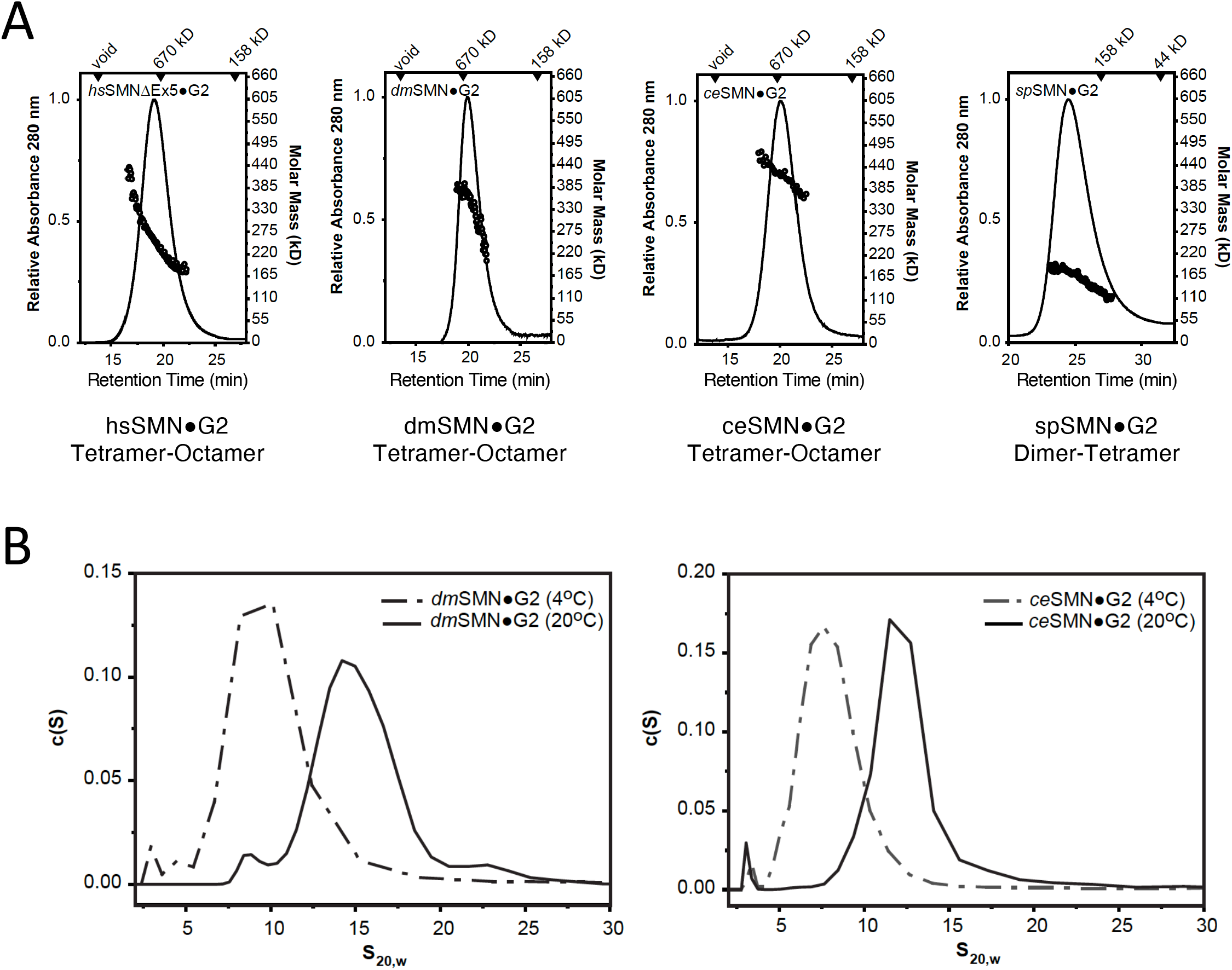
Biophysical properties of wild-type SMN ·Gemin2 complexes are conserved. ***A.*** SEC-MALS analysis performed at 20°C for *Homo sapiens* (*hs*), *Drosophila melanogaster* (*dm*), *Caenorhabditis elegans (ce)* and *Schizosaccharomyces pombe* (*sp*) SMN·G2 complexes. As previously observed for the human and fission yeast complexes [33, 34], SEC elution times suggest the presence of larger species in metazoans. However, *M*w values from the in-line light scattering indicate a range of oligomers in each case; the oligomeric range is assigned below each panel. ***B.*** Sedimentation velocity (SV-AUC) analyses of *d*mSMN·G2 (left) and *ce*SMN·G2 (right). At 25°C, the fruitfly complex sediments as a broad ~15 S peak but at 4°C a smaller ~8 S peak is observed. Similarly, the nematode complex sediments as a broad ~12S peak at 25°C but smaller ~7S peak at 4°C. See supplemental Fig. S2 for additional biophysical characterization of the *dm*SMN complex and its components.

The fly complex was unique among the systems examined in that both dmSMN and dmG2 could be purified separately and the complex reconstituted *in vitro*. In contrast, the human, worm, and yeast systems required bacterial co-expression to obtain soluble complexes. We used this opportunity to determine the properties of dmSMN and dmG2 alone. As anticipated, dmSMN is oligomeric, with a sedimentation coefficient of 11S, and dmG2 is largely monomeric, with a very modest dimerization constant detected by SE-AUC (Fig. S2). Previous studies with hsSMN and spSMN showed that the oligomeric behavior of the SMN·G2 complexes was recapitulated by fusions of the maltose binding protein (MBP) to the SMN YG boxes [33, 34]. This behavior is true of the fly complex as well; MBP-dmSMN(186-220) forms complexes spanning the tetramer-octamer range (Fig. S2). As a final test of the fruitfly proteins, we analyzed dmSMN·G2 complexes and MBP-dmSMN(186-220) fusions using size-exclusion chromatography in-line with synchrotron small-angle X-ray scattering (SEC-SAXS). The scattering profiles for the dmSMN·G2 complex were deconvoluted into two primary components: tetramers and octamers (Fig. S2). Thus, as analyzed by AUC, SEC-MALS, and SEC-SAXS at micromolar concentrations, metazoan SMN·G2 complexes exist as a mixture of oligomers spanning tetramers to octamers, whereas the fission yeast complex forms dimers and tetramers.

### Genetic and biophysical analysis of SMN mutations in S.pombe

Although *Smn* has been lost from the budding yeast genome, it is an essential gene in the fission yeast [39–41]. Fungal SMN proteins contain only two of the three conserved domains (Fig. 3A), the Gemin2 binding region and the YG box, both of which are essential for viability [41]. To understand the effects of YG box missense mutations on cell growth and SMN oligomerization potential, we carried out site-specific mutagenesis and compared phenotypes of the mutant proteins when expressed *in vivo* and *in vitro.* As summarized in Fig. 3, expression of the wild-type Smn1p construct fully complemented deletion of the endogenous *smn1* gene. In contrast, most of the mutant proteins failed to complement growth *in vivo* and were likewise affected in their ability to form SMN·G2 tetramers when expressed *in vitro* (Fig. 3). For example, replacement of three of the conserved ‘s-motif’ residues (Fig. 1A) with bulky residues like glutamine (S130Q, A134Q) or isoleucine (T138I) all caused defects in oligomerization and cell growth (Fig. 3). Substitution of other YG box alanines with glutamine (A141Q or A145Q) had little effect on growth or oligomerization.

**Figure 3:**
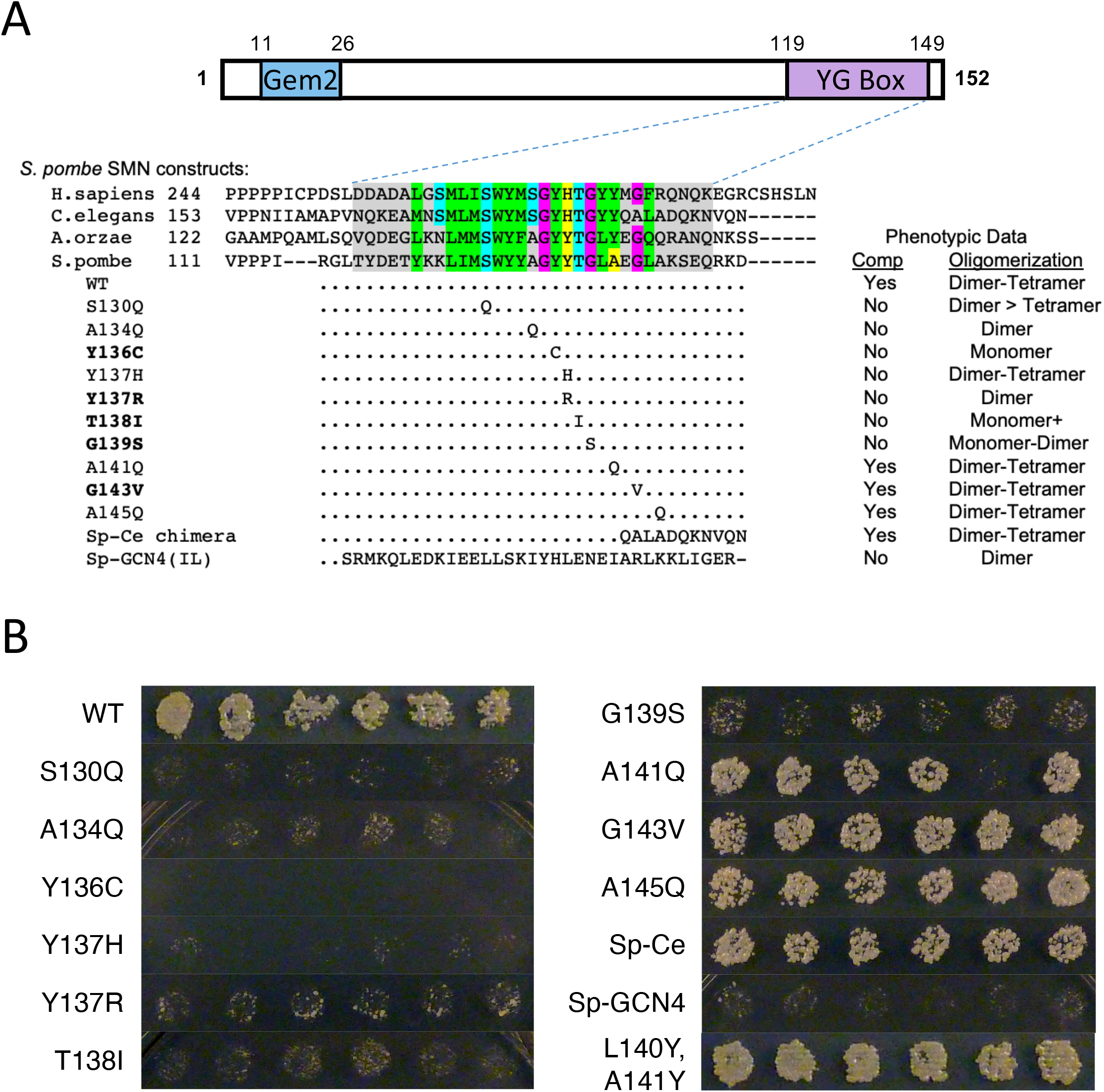
Genetic and biophysical characterization of fission yeast SMN mutants. ***A***. Cartoon of *sp*SMN protein, showing relative location of the Gemin2 (Gem2) binding domain and the YG box. An alignment of YG box sequences from the human (*H.sapiens*), nematode (*C.elegans*), köji mold (*A.orzae*) and fission yeast (*S.pombe*) SMN orthologs is shown for comparison. Genetic complementation analysis of a fission yeast *smn1* null allele was performed with either wild-type (WT) rescue construct or with a variety of chimeric or point substitution mutation constructs. Mutants that correspond to human SMA-causing missense alleles are shown in bold text. Ability to complement (Comp) the growth defect observed in the null mutant background is indicated. Recombinant SMN·G2 complexes of these same mutant constructs were generated *in vitro* and subjected to SEC-MALS and SE-AUC analysis, as described in Fig. 2. The range of oligomeric species detected by SEC-MALS is also indicated. See supplemental Table S2 for additional details regarding biophysical characterization of spSMN complexes. ***B.*** Complementation of *smn1+* deletion in *S.pombe.* Constructs expressing *smn1* variants under control of the *nmt1* promoter were transformed into haploid *S.pombe* containing a deletion of chromosomal *smn1+* and episomal *smn1* expressed from a *ura4+* plasmid. Individual transformants were patched onto minimal medium containing thiamine, then replica plated to dilution on selection plates containing 5-fluoroorotic acid (5-FOA). Strong growth on FOA medium requires loss of the *ura4+* plasmid and therefore complementation of *smn1Δ* by the *smn* construct.

As expected, modeling of human SMA patient-derived missense alleles in *S.pombe* revealed that mutations of highly conserved YG box residues (Y136C, Y137R, T138I and G139S) failed to complement the *smn1* deletion (Fig. 3). Moreover, all four of these mutations caused defects in SMN oligomerization to different extents (Fig. 3A). The spY136C protein is monomeric, consistent with studies of this mutation in other model systems that showed a failure to interact with itself in pulldown assays (e.g. hsY272C and dmY203C [23, 29]. We note that one of the SMA-causing alleles (spG143V) did not display a defect in complementation (Fig. 3A,B). Interestingly, overexpression of this same spG143V mutant protein in a wild-type *smn1* background was previously shown to impede growth of *S.pomb*e in a dominant negative fashion [40], but here we show that this allele is viable when expressed in the absence of the wild-type protein (Fig. 3B). Furthermore, and similar to previous findings in *Drosophila* S2 cells (dmG210V, [29]) and MBP-hsSMN(252-294) fusion constructs (hsG279V, [33]), we show that the spG143V construct is able to form higher-order oligomers (Fig. 3).

Given that this C-terminal glycine (hsG279) is the least well-conserved position among the three G-motif residues (see Fig. S1A), the spG143V result suggests that the observed metazoan SMA phenotype (human: Type I, [42]; fruitfly: Class 2, [43]) is caused by a distinct mechanism-of-action. Indeed, the C-terminus of the nematode SMN orthologue diverges in this region and a chimeric fusion of the yeast and worm YG boxes fully complements the *smn1* null mutation *in vivo* and displays an oligomerization profile that is very similar to that of the wild-type *S.pombe* protein *in vitro* (Fig. 3A). Thus, we conclude that the C-terminal G-motif glycine is not required for SMN self-interaction.

In order to determine whether dimerization alone is sufficient for SMN to carry out its functions in fission yeast, we generated a series of chimeric fusion constructs that replace the entire *S.pombe* YG box with a leucine zipper domain derived from the *S.cerevisiae* transcriptional activator, GCN4. The ~34 a.a. coiled-coil motif present within GCN4 has been thoroughly studied and can be tuned to generate dimers, trimers or higher-order oligomers [44]. When fused to spSMN, the GCN4-(IL) peptide (Sp-GCN4 chimera, Fig. 3A) forms obligate dimers when analyzed by SEC-MALS, SV-AUC, and SAXS (Fig. 3B and Table S2), whereas the GCN4-(LI) and -(II) fusion proteins form multiple higher-order oligomeric species (Table S2 and Fig. S6). Importantly, all three of these chimeric proteins fail to complement the *smn1* deletion *in vivo*. This result was not unexpected, as the YG box is thought to provide binding surfaces for Sm protein substrates as well as other members of the SMN complex [14, 27]. These data clearly show that multimerization, *per se*, is insufficient to support organismal viability.

### Oligomeric properties of human SMN·Gemin2 complexes bearing SMA-causing YG box point mutations

Thus far, we have only analyzed the oligomerization status of human SMA-causing YG box missense mutations in the context of MBP-fusion constructs [33]. To directly compare the molecular size data in Fig. 3B, we generated hsSMNΔ5oG2 complexes that contain SMN point mutations and analyzed their properties using SEC-MALS (Fig. 4). Exon 5 encodes a proline-rich region in hsSMN that causes the SMN·G2 complex to have reduced solubility, making biochemical analyses difficult. We therefore used the naturally occurring SMNΔ5 isoform for these experiments. As shown in Fig. 4A, hsSMN and hsSMNΔ5 contain all three conserved domains.

**Figure 4:**
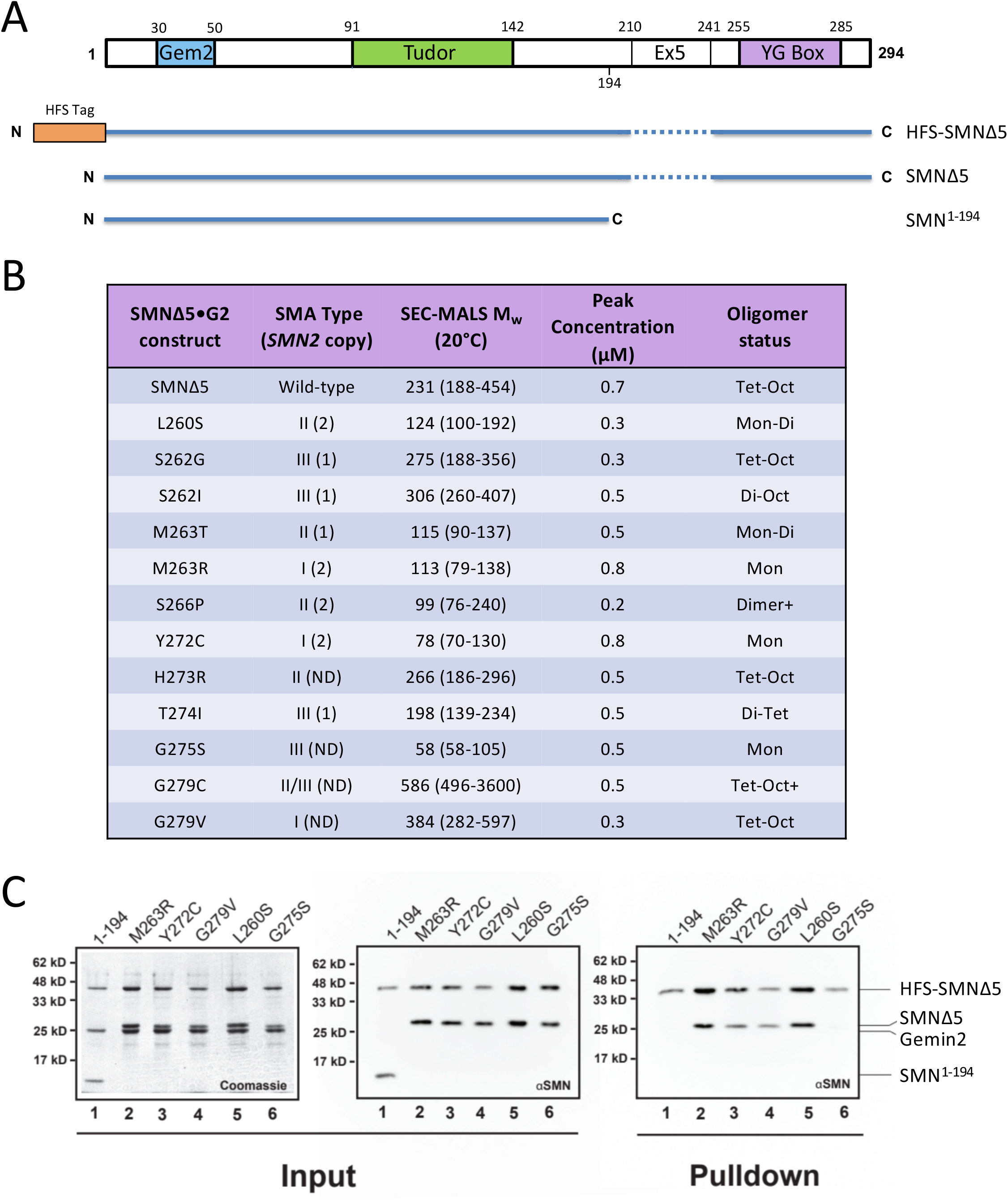
Biophysical characterization of human SMN·Gemin2 complexes bearing SMA-causing YG box missense mutations. ***A.*** Cartoon of *hs*SMN protein, showing the conserved YG box, Gemin2 binding (Gem2) and Tudor domains, along with location of exon5 (Ex5) sequences deleted in the SMNΔ5 construct. ***B.*** SEC-MALS analysis of SMA-causing point mutant constructs in the hsSMNΔ5 ·Gemin2 backbone. All of the mutations were generated on the SMNΔ5 backbone. Additional information regarding reported human *SMN2* copy number and SMA patient phenotype is also provided. ***C***. Formation of mixed oligomers between wild-type hsSMN and SMA patient mutations *in vitro.* A subset of the patient-derived mutations was screened for the ability to form mixed oligomers with wildtype SMNΔ5. The left two panels show Coomassie and western blot analyses of SDS-PAGE gels of the input material following bacterial co-expression and lysate clarification; the last panel shows a western blot of the resulting pulldown using Chitin-binding resin. As a negative control, the ability of a truncated SMN lacking the YG oligomerization domain (SMN^1-194^) was also assayed. After co-expression and elution from the chitin binding resin, four of the five patient mutant samples (M263R, Y272C, G279V, and L260S) demonstrated the ability to form mixed oligomers. Of the oligomeric constructs, only SMNΔ5(G275S) failed to co-purify with wild-type SMNΔ5.

For the most part, the severity of the SMA phenotype correlates well with the oligomerization potential of the corresponding SMN·G2 complexes, and with the previously reported oligomerization status of the corresponding MBP-fusion constructs (Fig. 4B; [33]). Several of the SMA point mutants are within a strongly conserved hydrophobic region that precedes theY, G, and s motifs (Fig. 1A). For example, L260S, M263T and M263R would be expected to weaken or disrupt the helical dimer shown in Fig. 1B and that is what we observe. Although the crystal structure of the hsSMN YG box [33] did not provide insights into much of this region, the longer N-terminal extension present in the spSMN YG box structure [34] indicates clear roles for these residues in extending the hydrophobic interface between helices.

In certain cases, the SMA-causing mutations have little effect on the oligomeric state of SMN·G2. Substitutions at human Ser262, His273, and Gly279 each result in complexes that can form higher-order oligomers. The hsG279V mutant is particularly interesting because one might expect that a bulky valine substitution would be incompatible with the close Gly-Gly contacts present in the YG box dimer. However, as discussed above for the equivalent yeast SMN substitution (spG143V), the dimer apparently tolerates a larger separation of helices near hsGly279, and the resulting structural perturbations do not affect formation of higher-order oligomers. Presumably, the structural changes associated with Cys and Val substitutions do interfere with some aspect of SMN biology, leading to the intermediate and severe SMA phenotypes observed.

The experiments summarized in Fig. 4B address the homo-oligomerization potential of SMA point mutations but do not indicate whether the mutant proteins are likely to form hetero-oligomers with wild-type SMN. To test this idea, we co-expressed a subset of the human YG box mutants with hexahistidine-FLAG-SUMO (HFS)-tagged SMNΔ5 and Gemin2, purified the complexes, and determined whether the untagged mutant proteins copurified with Ni-NTA beads (Fig. 4C). The M263R, Y272C, G279V, and L260S mutants can each interact with wild-type SMN. *In vivo*, these SMN variants would therefore be expected to display dominant negative phenotypes, consistent with the observations that dmM194R and dmY203C display incomplete dominance over the maternal SMN contribution in the fly [29] and that spY136C suppresses growth when co-expressed in fission yeast [40].

The hsG275S mutant does not interact with wild-type SMN (Fig. 4C). On a structural level, this is not a surprising finding, given that Gly275 is the central glycine in the YG box and even a single substitution in the interacting Gly-Gly pair in a dimer should be severely disruptive (Fig. 1B). This provides a compelling explanation for the milder SMA severity of this mutation, given that the serine substitution renders SMN monomeric. The relative lack of a dominant negative effect would also explain why the corresponding dmG206S mutant displays a slightly milder phenotype in the fly as compared to dmY203C [43].

### Genetic and biophysical analysis of YG box mutations in D.melanogaster

As shown in Fig. 5A, the overall structure of *Drosophila* SMN is similar to that of the human protein (Fig. 4A), and a homozygous null mutation of the endogenous *Smn* gene can be rescued by transgenic expression of a wild-type (WT) FLAG-tagged construct [45]. Previously, we showed that expression of disease-causing missense mutations in all three conserved SMN subdomains recapitulates the full spectrum of phenotypic severity observed in human SMA [29, 43]. For example, the *Drosophila* M194R, Y203C and G206S mutants are severe Class 1 alleles that die during larval stages, whereas Y208C, G210C/V (Class 2) and T205I (Class 3) display intermediate phenotypes that manifest during pupal or adult stages (Fig. 5A). Phenotypic comparison between SMA patient mutations and the corresponding animal model is complicated by human *SMN2* copy number variation [22]. In cases where the *SMN2* copy number is known, the severity observed in each fly mutant Class is well aligned with that of human SMA patient Type [43].

**Figure 5:**
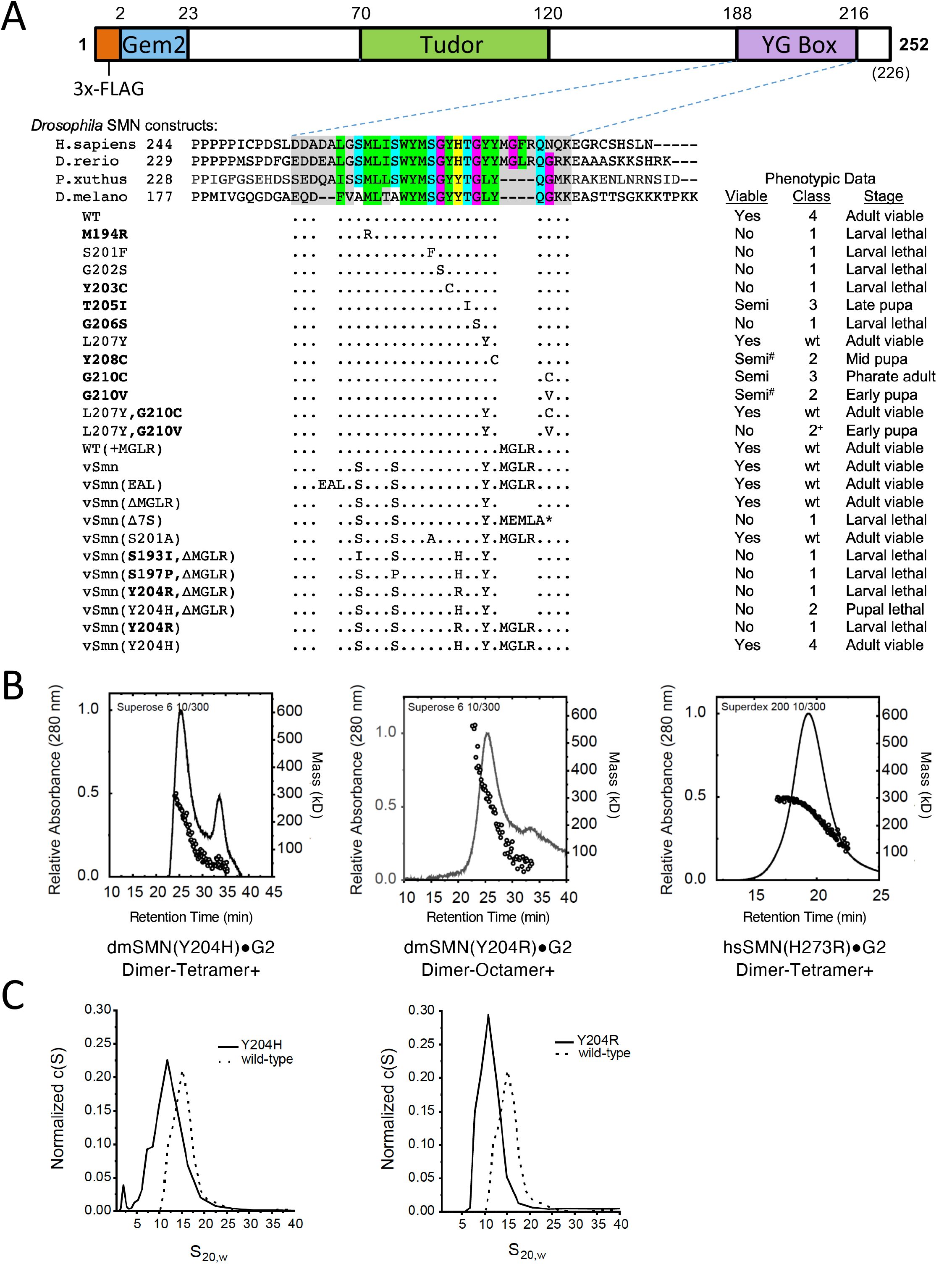
Genetic and biophysical characterization of *Smn* YG box mutations in *Drosophila.* *A.* Cartoon of the *dm*SMN protein, showing the conserved YG box, Gemin2 binding (Gem2) and Tudor domains, along with the location of the 3x-FLAG tag used for transgenic rescue experiments. An alignment of YG box sequences from the human (*H.sapiens*), zebrafish (*D.rerio*), butterfly (*P.xuthus*) and fruitfly (*D.melanogaster*) SMN orthologs is shown for reference. Phenotypic comparisons of an extensive panel of *Drosophila* YG box substitution mutations are summarized below the alignment. With the exception of S201F and G202S, which are point mutations in the endogenous *Smn* gene [56], the rest of the panel is comprised of transgenic constructs that have been recombined with an *Smn* null allele [29]; this work). Genetic complementation analysis was performed using either a wild-type (WT) or mutant *Flag-Smn* transgene. Mutants that correspond to human SMA-causing missense alleles are shown in **bold** text; the classification system used for fly SMA models was described previously [43]. See Fig. S4 for pupal and adult viability analysis of the fifteen new transgenic fly lines used in this work. Mutant lines that eclose at low frequency (adult escapers) are considered semi-viable; those marked with a # sign are incapable of establishing an independent breeding colony. Class 2^ denotes animals that pupate but display a significant advancement in lethal phase or reduction in eclosion frequency. * = stop codon. ***B***. SEC-MALS analysis of SMN·G2 complexes containing mutations at hsH273 (H273R) or dmY204 (Y204H and Y204R) aimed at modeling an SMA-causing missense allele in human *SMN1. **C**.* Analytical ultracentrifugation of *Drosophila* SMN·G2 complexes. c(S) sedimentation distributions of wild-type (WT) or mutant (Y204H and Y204R) are shown. Note, additional biophysical analyses of other *Drosophila* SMN constructs are also presented in Table 1 and Figs. S2 and S7.

Phenotypic differences are illuminating. For instance, we note that the Tudor domain mutants hsI116F/dmI93F and hsW92S/dmF70S are Type I SMA mutations in humans [46, 47] but display much milder Class 3 phenotypes in flies [29]. Interestingly, we recently found that these two fly mutations are highly temperature-sensitive and become differentially more severe (Class 2^) when animals are reared at elevated temperatures closer to those of mammals [48]. Among the few discordant YG box mutants, the hsG275S (Type III) and dmG206S (Class 1) pair is interesting because, as discussed above, the hsG275S complex is monomeric (Table 1). Although *SMN2* copy number was not determined in the single reported patient bearing this *SMN1* mutation [49], one would have expected a much stronger phenotype. We therefore generated the corresponding SMN·G2 complex (dmG206S) and found that it too is primarily monomeric, as measured by SEC-MALS. Thus, the biophysical properties of recombinant dmG206S (Table 1) are consistent with those of hsG275S as well as with results of GST-pulldown assays in *Drosophila* S2 cells [29].

**Table 1:**
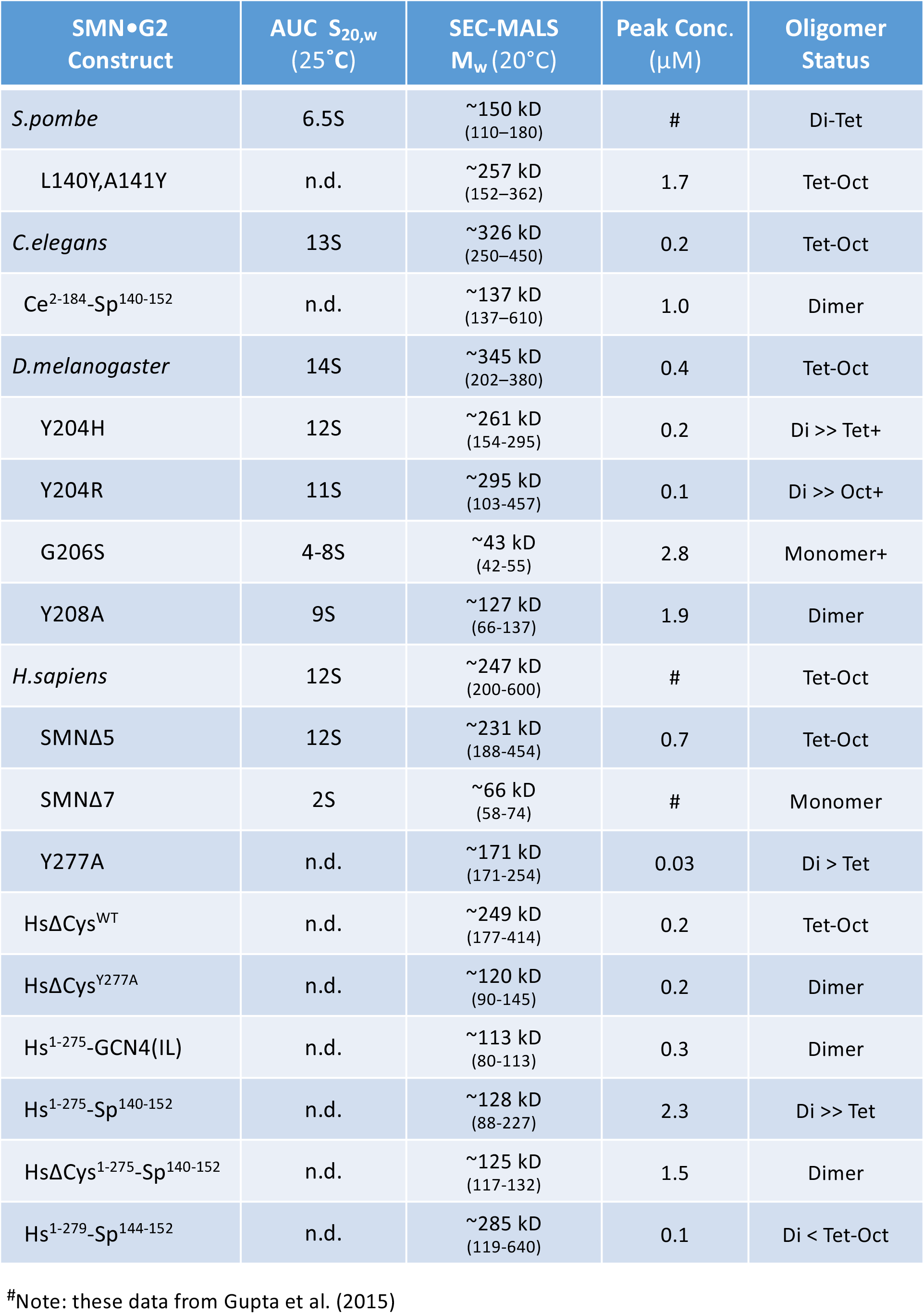
Oligomeric Properties of SMN·G2 Variants

Another instance where unknown *SMN2* copy number hinders interpretation of the SMA phenotype is at Gly279. Mutation of this residue to Cys (hsG279C) results in mild Type II/III SMA [50], whereas a Val substitution (hsG279V) causes severe Type I SMA [42]. In flies, the corresponding mutations are dmG210C and dmG210V, which were designated as Class 3 and Class 2 alleles, respectively [43]. As mentioned above (Fig. 3), the fission yeast spG143V mutant is viable. The model in Fig. 1B shows that human Gly279 is tightly packed against Tyr276, whereas in flies and yeast the tyrosine is replaced by leucine. In the fly model, Gly210 is tucked into a hydrophobic pocket created by Leu207 and the aliphatic side chain of Lys211 (Fig. 1B), and so leucine is predicted to be more tolerant of a valine substitution at Gly210 than is tyrosine. To test this prediction, we generated single and double substitutions at Leu207 and Gly210 in the fly. As expected, the dmL207Y controls are fully viable, whereas a small, but reproducible, fraction of dmG210V mutants can complete development (Fig. 5A; [43]). By contrast, the dmL207Y,G210V double mutants display a phenotype that is more severe than dmG210V alone and are completely inviable, with no eclosing adults (Fig. S4A). On the other hand, the dmL207Y-G210C animals display a phenotype that is less severe than dmG210C alone and are viable (Figs. 5A, S5A). The results show that Leu207 is indeed more tolerant of a G210V substitution than is Tyr207.

As an animal model, *Drosophila* arguably provides a better overall indication of SMN activity *in vivo* because mutant proteins can be expressed and analyzed in the absence of wild-type SMN [43]. However, due to sequence differences between the human and fruitfly YG box, certain SMA-causing point mutations could not be effectively modeled using the WT *Drosophila* backbone (Fig. 5A). Phylogenetic comparison of the YG box domains from nine different vertebrate clades (see Fig. S3A) suggests that there has been a deletion of four residues in the C-terminal region of the fruitfly YG box relative to the human sequence (residues 278MGFR281). Among these four residues, Phe280 is a clear phylogenetic outlier as this residue is typically a leucine (Fig. S3A). We therefore inserted a sequence encoding MGLR at the corresponding position in fly SMN to create the WT(+MGLR) strain. As shown in Fig. 5A, this allele is fully viable. Indeed, these animals eclose at slightly higher frequencies than do those of the WT rescue strain.

In addition, the N-terminal half of the human YG box contains a few other sequence differences, notably at conserved s-motif or Y-motif positions (Fig. S3A). Therefore, we generated additional vertebrate-like transgenic rescue lines (vSmn, vSmn^EAL^, and vSmn^ΔMGLR^) to serve as baseline controls. As summarized in Fig. 5A, each of these three fly strains is fully viable, demonstrating that neither the insertion of the MGLR residues nor substitution of Ser for Ala, or Leu for Phe at the other positions has any negative effects on organismal viability. Once again, these *vSmn* strains are healthier overall than the WT rescue line (see Fig. S4 for details). Also as expected, truncation of the fly protein in this region with MEMLA* to model SMNΔ7, the predominant human *SMN2* gene product, caused early larval lethality (Fig. 5A, vSmnΔ7S).

However, when we attempted to model one particular SMA patient-derived mutation (hsH273R, Fig. 4B), substitution of the corresponding human wild-type residue into the fly protein (vSmn^Y204H,ΔMGLR^) resulted in complete pupal lethality (Figs. 5A, S5B). Mutation of this residue to model the disease-causing allele (vSmn^Y204R,ΔMGLR^) significantly worsened the phenotype (Figs. 5A, S5B). Consistent with these findings, fission yeast spY137H and Y137R mutants failed to complement and displayed an impaired growth defect (Fig. 3A). Interestingly, spY137R was entirely defective in formation of higher-order SMN oligomers (Fig. 3A), whereas hsH273R was only partially impaired (Fig 4B). We therefore generated dmY204H and dmY204R constructs and analyzed their solution properties in complexes with Gemin2, as described above. As shown in Fig. 5B, these constructs both show a significant shift towards dimers and a decrease in higher molecular weight species. The trace for dmY204H is biphasic, with a clear peak in the monomer-dimer range (Fig. 5B), whereas the mass profile for dmY204R is consistent with the presence of monomers to large aggregates.

As described previously, hsHis273 sits on the outer side of the helix and makes no direct contacts along the dimer interface (see Fig. 1B). So why is the dmY204H/spY137H substitution toxic in flies and yeast? As shown in the yeast crystal structure (Fig. S1B) and the fly model (Fig. 1B), Tyr204 packs against the adjacent Tyr203 residue and helps to buttress the Y-G interaction that forms between Tyr203 and Gly206 on the other strand of the dimer. Histidine is polar and does not make the same hydrophobic contacts, and thus may not function as well in this role (see Discussion). The dmY204R substitution is even more polar and destabilizing, affecting both dimerization as well as formation of higher-order oligomers (Fig. 5B). Interestingly, insertion of the vertebrate MGLR helix extension completely suppresses the toxicity of the histidine substitution (Fig. 5A, compare vSmn^Y204H^ to vSmn^Y204H,ΔMGLR^), whereas the SMA-causing arginine substitution remains inviable (compare vSmn^Y204H^ to vSmn^Y204R^). We conclude that the fruitfly YG box is unable to support a histidine at Tyr204, affecting both higher-order multimerization as well as dimerization. In the human system, the hsH273R mutation does not appear to interfere with dimerization; rather, it drives the equilibrium towards lower-order multimers (Fig. 5B).

### Formation of higher-order SMN oligomers is required for metazoan viability

Our detailed structure-function analysis of the YG box has shown that mutations which reduce the oligomerization potential of SMN *in vitro* also tend to have a negative impact on organismal viability *in vivo* (Figs. 3-5). However, none of the SMA patient-derived missense mutations analyzed thus far allow dimer formation but prevent assembly of higher-order multimers. We therefore sought to identify a missense mutation that causes metazoan SMN to form obligate dimers *in vitro* and then determine its phenotype in an animal model. Toward that end, we focused on trying to understand the aforementioned discrepancy between the oligomerization status of wild-type fungal and metazoan SMN·G2 complexes.

As summarized in Table 1, animal SMN·G2 complexes mainly form tetramers and octamers at micromolar concentrations *in vitro*, whereas those of fission yeast exist primarily as dimers along with some tetramers. Chimeric constructs with yeast N-termini and human C-termini (Sp^1-117^-Hs^253-294^) form tetramers and octamers. To better understand this phenomenon, we generated reverse chimeras with C-terminal portions of yeast SMN fused to either human (Hs^1-275^-Sp^140-152^) or nematode (Ce^2-184^-Sp^140-152^) N-terminal regions. Interestingly, these complexes were primarily dimeric, although some tetramers were also detected (Table 1). A fusion bearing only the distal C-terminus of the yeast protein (Hs^1-279^-Sp^144-152^) restored the ability of the human construct to form octamers (Table 1) and allowed us to map the residues that were largely responsible for this effect to the region encompassing *S.pombe* residues 140LAEGL145 (see Fig. 6A). Thus, we conclude that the C-terminal portion of the fission yeast YG box contains determinants that limit formation of higher-order oligomers.

**Figure 6:**
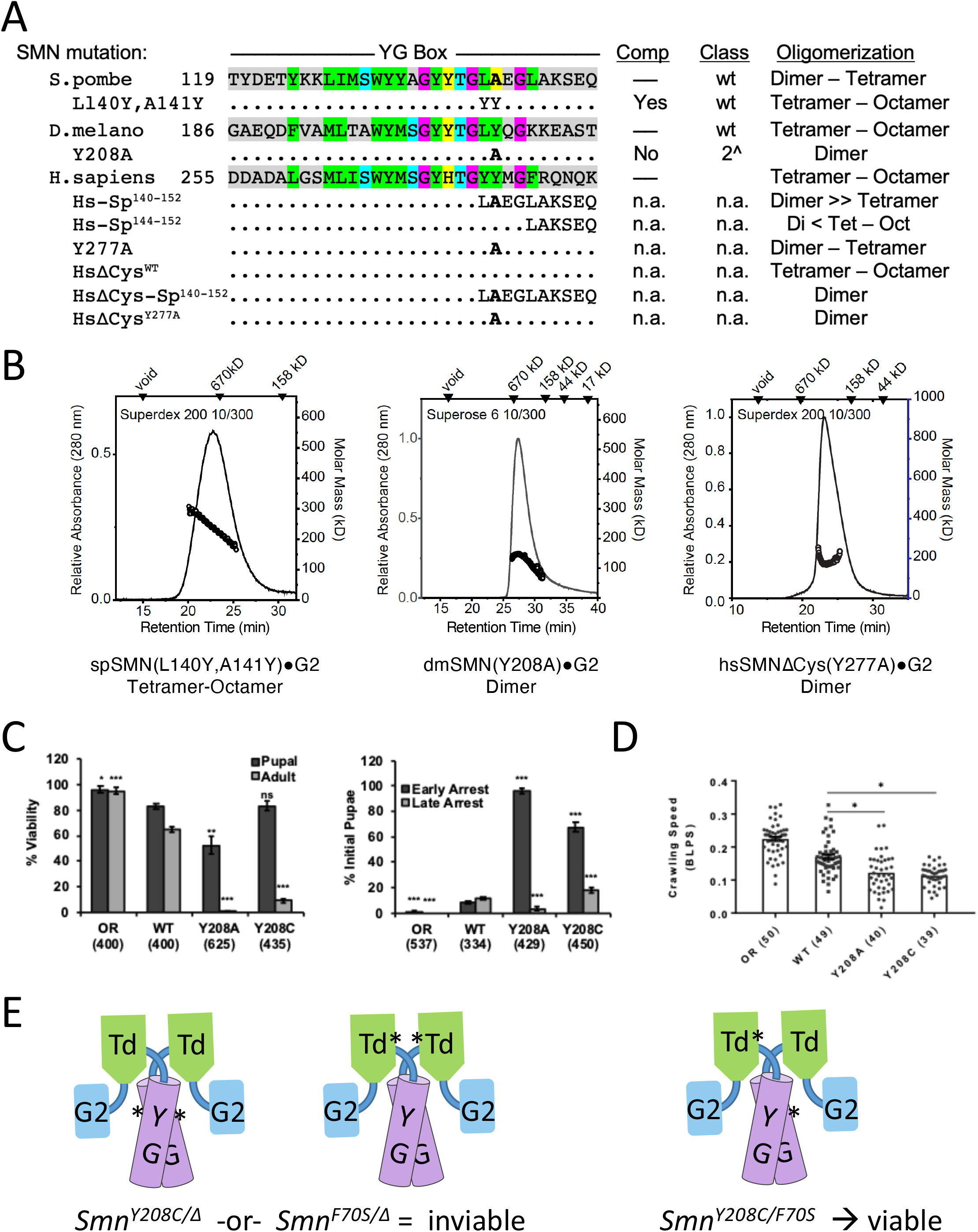
Identification of a specific YG box residue critical for formation of higher-order SMN multimers. A structure-function analysis was carried out in parallel using fission yeast, fruitfly and human SMN. ***A.*** Genetic complementation (Comp) and biophysical analyses (Oligomerization) of C-terminal chimeras and point substitution mutants. ***B***. SEC-MALS analysis of SMN·G2 complexes of yeast spSMN^L140Y,A141Y^, dmSMN^Y208A^ and hsSMN^Y277A^. ***C.*** Developmental viability of control Oregon Red (OR) flies or animals expressing transgenic Flag-tagged *Smn* wild-type (WT), Y208C or Y208A rescue constructs in the background of an *Smn* null mutation. Left panel: % Viability is the proportion of animals that survive to the pupal (darker gray) or adult (lighter gray) stages, relative to the number of larvae initially collected (n-values in parentheses). Right panel: Breakdown of fraction of animals that arrest during early vs. late stages of pupal development. ***D***. Larval locomotion analysis. Average crawling speed, measured in body lengths/sec, in wandering third instar larvae. Data points correspond to measurements of individual larvae. n-values for each genotype are shown in parentheses. Statistical analysis used in panels *A-D*: Asterisks above the data bars indicate significance vs. WT rescue line from one-way ANOVA using the Dunnet correction for multiple comparisons. *p < 0.05, **p < 0.01, ***p < 0.001. p > 0.05 is not significant (ns). ***E.*** Cartoon of intragenic complementation analysis, illustrating how Tudor and YG box mutations can complement each other. For additional details, see [29] and supplemental Fig. S5.

Inspection of the *S.pombe* YG box shows that it is a bit of a sequence outlier, even when compared to other fungal SMN orthologs. Indeed, the fungal consensus in this region is 140LYEGQ145 (numbering per spSMN, see Fig. S3B). Although substitutions are clearly allowed, *S.pombe* is the only species among the nine clades we surveyed with an alanine at position 141 (see Fig. S3B). To better model the human sequence, we generated an spL140Y,A141Y mutant construct and tested it for complementation in an *smn1* deletion background, as described above. As shown in Fig. 3B, this allele fully rescued the growth defect of the null allele *in vivo.* When assayed *in vitro* for oligomerization using SEC-MALS, we found that it displayed a tetramer-octamer distribution (Fig. 6B) like that of human, fruitfly and nematode SMN·G2 complexes. Furthermore, we note that in flies, the dmL207Y mutation (spL140Y) also improved the viability phenotype as compared to the WT Flag-*Smn* rescue transgene, whereas expression of an SMA-causing mutation at the adjacent residue, hsY277C/dmY208C, significantly impaired viability (Fig. 5A). Collectively, these findings implicate spAla141 as an anti-oligomerization determinant.

We therefore generated SMN·G2 constructs bearing the corresponding *Drosophila* and human mutations in SMN and determined their oligomerization status using SEC-MALS. As shown in Fig. 6B, the dmY208A complex is strictly dimeric, whereas the data for the hsY277A complex are consistent with a dimer-tetramer equilibrium. As a control, we generated a Hs^1-275^-GCN4(IL) chimeric fusion and again found that, in complex with Gemin2, this construct forms obligate dimers (Table 1). We hypothesized that sequences outside of the YG box might contribute to the modest differences observed between the dmY208A and hsY277A constructs. Conspicuously, Dreyfuss and colleagues previously identified two poorly conserved cysteine residues in the human protein (Cys60 and Cys250, see Fig. S3A) that can form SMN-SMN disulfide cross-linkages following treatment with various oxidizing agents [51].

Among the eight cysteines in hsSMN, three of them are located within the Tudor domain and are very highly conserved. We therefore generated a version of SMNΔ5 containing Ala substitutions at each of the other five Cys residues (Cys 60, 146, 231, 250 and 289). This construct (HsΔCys^WT^, see Table 1) was co-expressed and purified together with Gemin2 as described above, and then analyzed by SEC-MALS. As shown in Fig. 6A, this complex is best described by a tetramer-octamer equilibrium distribution, like that of the control SMNΔ5oG2 complex. Importantly, when generated in the ΔCys background, both the HsΔCys-Sp^140-152^ chimera and the HsΔCys^Y277A^ mutant form strict dimers with no evidence of tetramer formation (Fig. 6B, Table 1). Thus, when evolutionarily variant residues are considered, the SMN·G2 oligomerization results are completely concordant across multiple eukaryotic phyla, including *Chordata*, *Arthropoda*, *Nematoda* and *Ascomycota* (Table 1).

Having established the importance of *Drosophila* Tyr208 to the formation of higher-order SMN oligomers *in vitro*, we generated transgenic flies bearing a dmY208A mutation to study its effects *in vivo*. We assayed organismal viability, lethal stage and larval locomotion phenotypes in comparison to the previously described SMA missense mutation strain, dmY208C [43], as well as to wild-type controls. As shown in Fig. 6C, control animals complete development and eclose at high frequency, unlike the two missense mutants. However, dmY208A animals display a significantly more severe phenotype than do the dmY208C mutants, as a small fraction of the latter reach adulthood. By contrast, none of the dmY208A mutants eclose as adults, and they are also significantly impaired during pupariation as compared to either the dmY208C or the WT control line (Fig. 6C). Both missense mutations display early onset SMA-like phenotypes, appearing during larval stages. Locomotion analysis of wandering third instar larvae shows that crawling velocity of both mutants is significantly reduced (Fig. 6D). Finally, intragenic complementation analysis of SMA-causing mutations within the YG box versus the Tudor domain suggest that these two domains perform independent functions (Figs. 6E and S5).

In summary, these experiments show that animals expressing a YG box mutation that allows dimerization but prevents assembly of higher-order SMN oligomers is incompatible with metazoan life. The results also call into question whether unicellular organisms like fission yeast require formation of SMN·G2 oligomers larger than tetramers to support cell growth.

### A structural model for SMN tetramer formation

The combined data from three different model systems point to the relative importance of four distinct YG box residues in the assembly of higher-order SMN multimers. Obviously, hsY277/dmY208/spA141 is one of those residues, as detailed above (Figs. 5 and 6). In addition, we showed that mutations at hsH273/dmY204/spY137 shift the oligomeric equilibrium towards lower-order species (Figs. 3, 4 and 5). The other two residues are hsS266/dmA193/spS130 and hsS270/dmS201/spA134, as mutations at these positions are inviable and cause SMN to form dimers in fission yeast (Fig. 3).

Among the lesser studied residues within the SMN YG box are those that comprise the S (serine)-motif [33], which for reasons that will become apparent, we have renamed as the s (small)-motif (Fig. 1A). Due to the presence of conserved serine/threonine residues, especially within higher eukaryotic organisms, we and others initially assumed that phosphoregulation of these residues was somehow important for SMN biology. Indeed, mutation of hsS270/dmS201 to either aspartate or alanine has dramatic (and opposite) effects on SMNΔ7 protein stability in both human and *Drosophila* cells [52, 53]. Notably, this residue forms a key part of an E3 ubiquitin ligase phosphodegron [52], but the serine itself is not essential for viability in the context of full-length SMN, as dmS201A mutants are viable and fertile (Fig. 5A). By contrast, substitution of a bulky Phe residue at this position renders the dmS201F protein unstable *in vivo* [54, 55], and the animals are inviable (Fig. 5A; [56].

Glutamine scanning mutagenesis of various small residues in the yeast YG box (Fig. 3A) reveals that insertion of a bulky and polar Gln residue can be tolerated at certain positions in spSMN (e.g. spA145Q and spA141Q), whereas at other positions (e.g. spS130Q and spA134Q) it cannot. Quizzically, the residues that are intolerant to Gln substitution tend to be located at s-motif positions that do not lie along the direct YG zipper dimerization interface (Fig. 1A,B). To better visualize this information, we generated a helical wheel diagram of the human YG box dimer. As illustrated in Fig. 7A, the Tyr and Gly residues that are directly involved in SMN dimer formation are located along the same face of the alpha helix, whereas all four of the residues implicated in higher-order multimerization are on the opposite side. Furthermore, the conserved small residues (Ser266, Ser270, and Thr274) are in perfect register, suggesting the presence of a novel binding interface as shown by orthogonal views of the YG-box dimer in Fig. 7B.

**Figure 7:**
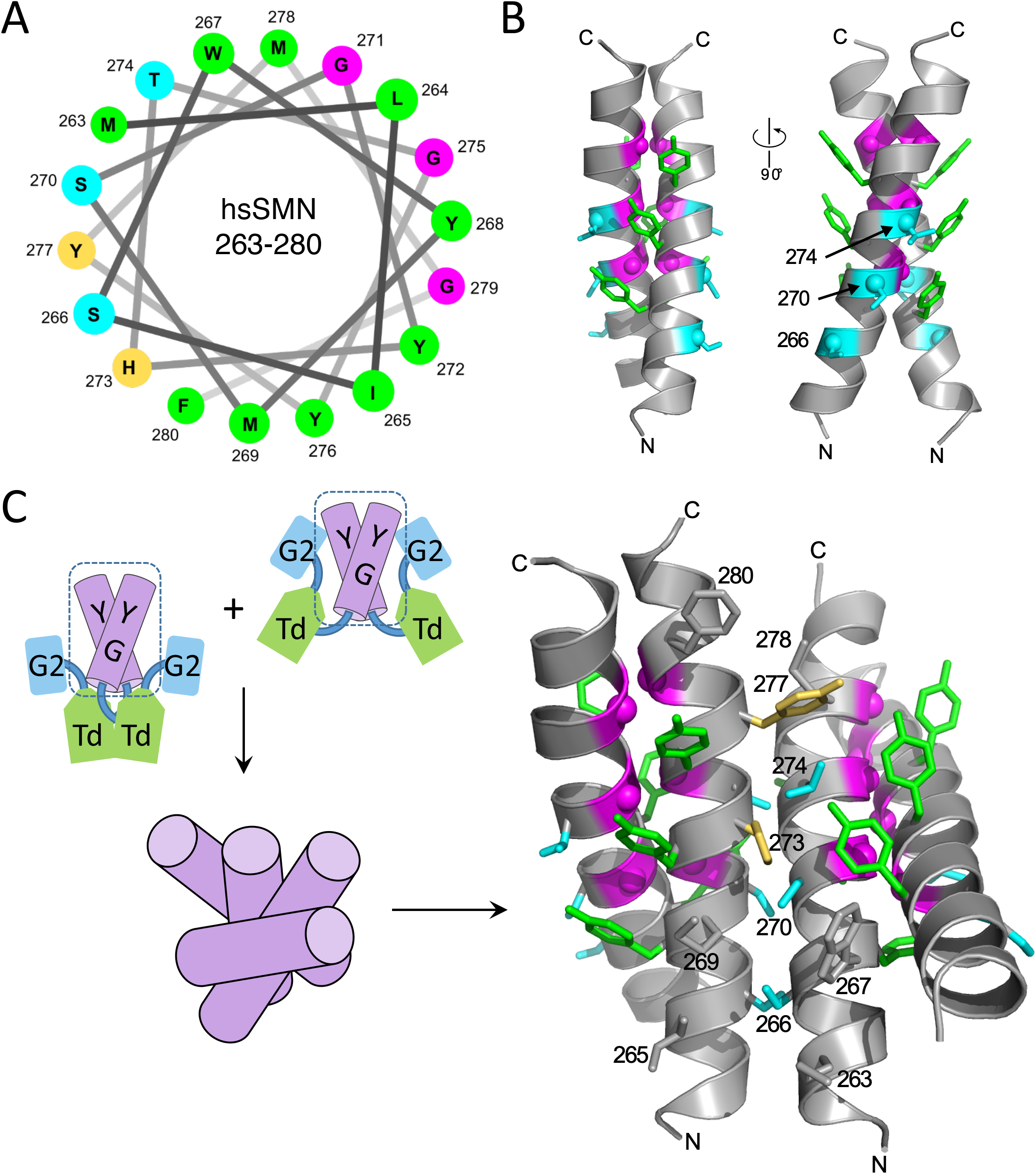
Molecular modeling of SMN tetramer formation. ***A.*** Helical wheel diagram of an *hs*SMN YG box. Color coding of residues is per Fig. 1A. Residues critically involved in forming the Tyr-Gly (YG) dimer interface are located on one side of the wheel. Conserved small residues (S266, S270, and T274) as well as key residues implicated in higher order multimer formation (H273 and Y277) are located on the opposite side of the wheel. ***B***. Orthogonal views of the human YG-box dimer illustrating the location of the “s-motif” opposite to that of the YG interface. ***C***. Proposed model of a YG-box tetramer. The s-motif residues, along with His273 and Tyr277, form a novel helix-helix interface. Helical crossing is right-handed, as found in other membrane proteins that employ Ala, Ser, Thr, and Gly residues to form similar interfaces. His273 and Tyr277 are expected to play key roles in stabilizing tetramers and higher order oligomers in this model.

Tetramers can form by two distinct mechanisms: one in which all four subunits are equivalent (symmetric bundle), and one that involves self-association of dimeric subunits (dimer of dimers). In a symmetrically bundled tetramer, the same residues are involved in forming the intersubunit contacts of both the dimeric and tetrameric forms, whereas in the dimer of dimers model, different binding surfaces are used for dimerization versus tetramerization. Previously, we carried out cross-linking studies with pre-formed SMN dimers and measured subunit exchange after mixing tagged and untagged complexes. The results showed that, *in vitro*, spSMN forms tetramers via the association of stable dimers [34]. As summarized in Fig. 7A-B, our *in vivo* and *in vitro* mutagenesis experiments not only support this notion, but clearly define a second intersubunit binding surface along the opposite face of the YG helix.

We therefore carried out an *in silico* analysis to consider which tetramer models would be most consistent with the available experimental data. We first asked whether the dimers might associate in a parallel vs. anti-parallel fashion. Two SAXS approaches using SMN YG box constructs support a parallel dimer-of-dimers model. In the first experiment, we compared SAXS data for dimeric and tetrameric spSMN·G2 complexes (Fig. S7). The radii of gyration (Rg) of the dimer and tetramer can be used to estimate the distance separating the two dimer components within the tetramer by application of the parallel axis theorem [57]. This distance is ~84 Å, a value that is only slightly larger than the 60 Å length of the SMN YG box dimer. Given that the spatial extents of the dimer (230 Å) and tetramer (270 Å) indicated by SAXS are quite large, it is unlikely that the centers of mass of the two dimers are located near the ends of the YG boxes, as would occur in an antiparallel tetramer (see Fig. S7). An antiparallel arrangement also implies a larger increase in complex dimensions than what is actually observed in the tetramer.

In the second experiment, we used the SAXS profile of tetrameric MBP-hsSMN(252-294) to compare models of parallel vs. antiparallel arrangements of YG box tetramers *in silico* (Fig S8). In this approach, the YG box components were fixed and the MBP domains were allowed to sample an ensemble of non-overlapping positions via a flexible SMN(252-255) linker, using the program CORAL [58]. There was good agreement between the experimental SAXS data and the predicted scattering of a parallel tetramer (χ^2^=1.4), but poor agreement with that of the antiparallel tetramer (χ^2^=5.3). We therefore predict that SMN YG box dimers associate in a parallel fashion to form tetramers.

To identify a plausible model for the parallel association of SMN dimers, we searched for examples of helix-helix interactions involving YG box s-motif residues (Figs. 1A, S1A) such as serine, alanine, and threonine. Helix-helix packing that features small residues at the interface is common among membrane proteins, where the interfaces are enriched for Ala, Ser, Thr, and Gly residues, and the helix crossing is generally right-handed [59]. Indeed, an extreme example of this type of helix-helix interaction is the glycine zipper, where Gly residues mediate intimate contact between the helical backbones as occurs in the SMN YG-box dimer. In contrast, the well-studied coiled-coil interface is enriched in Leu, Ile, and Val residues and the crossing is left-handed [60, 61].

We therefore generated a model for SMN tetramerization based on previously observed small residue interfaces, e.g. from the glycerol facilitator protein [62]. Superimposing one helix from each of two YG box dimers onto a templated helical pair from pdb entry 1FX8 [62], the resulting model (Fig. 7C) places the conserved s-motif residues, along with His273 and Tyr277, at the interface between SMN dimers. Substitutions at these positions would be expected to interfere with tetramer stability. Closely related alternative models may also be plausible, but as discussed below, the arrangement shown in Fig 7C provides a useful structural platform for understanding the mechanism of SMN oligomerization.

## Discussion

Transmembrane proteins frequently contain GxxxG motifs that promote dimerization [61]. This motif is often extended to a (G,A,S)xxxGxxx(G,S,T) sequence, known as a glycine zipper [32], which is thought to mediate oligomerization. Although helices bearing these motifs interact with distinct right-handed crossing angles, glycine zipper proteins form symmetrically bundled oligomers that associate in a front-to-back fashion, whereas GxxxG dimers employ face-to-face packing [32, 63]. The SMN YG box contains an extended GxxxGxxxG motif, however the Gly residues are face-to-face as they are in the GxxxG dimers (Fig. 1B). This unique feature of the YG zipper suggests that once an SMN dimer forms, that same binding surface should no longer be available to generate oligomers. Our experimental data are not only consistent with this model, but they identify specific YG box residues that mediate formation of higher-order multimers.

### Structural aspects of SMN oligomerization

The residues present within the tetrameric interface in Fig. 7C are not normally found in the interfacial positions of soluble protein coiled-coils, which are dominated by Leu, Ile, and Val [60]. In SMN, the core interfaces involved in forming both dimers and oligomers utilize small residues like Ser, Ala, Thr and Gly. In the model, we suggest that Ser266 and Ser270 are both largely buried in the interface of an alternative, right-handed helix-helix interaction where they could form inter-helical hydrogen bonds with the polypeptide backbone, as observed in high resolution structures of membrane proteins ([59]; see above). This positioning would explain the sensitivity of the corresponding spSMN and dmSMN residues to substitution by larger side chains, but not by smaller ones (Figs. 3 & 5). The model also explains how formation of higher-order multimers could protect SMN from degradation via the Ser270 phosphodegron [52], which is solvent exposed in the dimer.

His273 and Tyr277 play more standard roles at the tetramer interface, where their “knobs” pack into the “holes” generated by side chains on the partner helices. Both would be expected to play important roles in tetramer formation in this model, where the bulky His and Tyr side chains are predicted to make substantial interactions. Evolutionarily speaking, His273 is usually a Tyr in the SMN proteins of lower organisms, with Gln present in a very small number of cases (Figs. 1A, S1A). A large hydrophobic residue is even more strongly conserved at Tyr277, where the Ala found in *S.pombe* is an outlier, as noted earlier. The weakened stability of spSMN tetramers compared to those of the metazoan orthologs we studied can be readily explained by the lack of a bulky hydrophobic residue at Ala141, the position corresponding to Tyr277 of hsSMN. The same argument explains the loss of higher-order oligomerization that we found for the hsY277A and dmY208A mutants (Fig. 6, Table 1). Indeed, the strong temperature dependence of higher-order oligomerization observed in the metazoan SMN complexes (Fig. 2) also suggests a hydrophobic driving force. The defective phenotypes of hsY277C and dmY208C are also consistent with the idea that the residue at hsTyr277 plays a key role in mediating higher-order multimerization and that oligomers larger than SMN dimers are required for proper animal development.

In addition to the four residues discussed above, additional SMN YG box residues contribute to the interface shown in Fig. 7C. Met263, Trp267, Met269, Thr274, Tyr276 and Met278 all provide flanking hydrophobic interactions and Phe280 is positioned at the C-terminal end of the helix-helix interface. There is a strong correlation between the presence of His at position 273 (hsSMN numbering) and a large hydrophobic at 280 (see Figs. S1A and S3). When Tyr is present at position 273, the residue at 280 is more variable and is often not hydrophobic. This finding suggests that His273 might be less effective than Tyr in this position but is compensated by a strong hydrophobic residue at position 280, and most often a flanking hydrophobic at 278. These observations provide a plausible explanation for the lethal phenotype and weakened tetramerization of the Y204H substitution in dmSMN, where the residues corresponding to Met278 and Phe280 in hsSMN are Gln and Lys, respectively (Table 1 & Fig. 5). Insertion of sequences that place Met and Leu in these positions (+MGLR) results in a more “histidine friendly” environment and a viable phenotype (Fig. 5).

The crucial role of His or Tyr at position 273 in our model also helps to explain the sensitivity of these residues to mutation. The human H273R substitution causes Type II SMA [64], and the Y204R mutation is Class 2 in flies (Fig. 5A). The corresponding Y137R substitution in fission yeast fails to support growth (Fig. 3). Although the hsSMN H273R variant does not display a defect in oligomerization (Fig. 4B), the flanking hydrophobics and the presence of Phe280 presumably compensate for the Arg substitution. In the absence of such compensatory residues, both dmY204R and spY137R mutants are primarily dimeric (Fig. 3, Table 1). In summary, a model wherein two YG-box dimers associate via a righthanded small residue interface provides a plausible mechanism for formation of higher-order oligomers of SMN.

### Regulation of SMN multimerization

The model we outline above explains a large body of genetic, phylogenetic and biochemical data, and provides an obvious mechanism for forming larger oligomers, because the tetramer has two binding surfaces available for interaction with other SMN dimers or tetramers. What factors limit the extent of SMN polymerization? *In vitro*, SMN·Gemin2 complexes in the nM-μM range occupy multiple oligomeric states, and their hydrodynamic properties show that they adopt highly extended (non-globular) conformations. Clearly, the N-terminal portion of SMN is an important determinant of its oligomerization potential and overall solubility, as YG box constructs tagged with small epitopes or synthetic YG box peptides form large, insoluble aggregates. By analyzing a series of MBP-YG box fusions, we found that we could control the size of the complexes formed (n = 1, 2, 4 or 8) by very small changes in the length of the linker (Ref. [33] and this work). As illustrated in Fig. 7C, the amino terminal region of SMN could also adopt different conformations that might limit the overall number of multimers that form, with analogy to a bouquet of flowers in a vase. Thus, steric factors including N-terminal composition and linker length are critical determinants in regulating the extent of YG box oligomerization.

*In vivo*, post-translational modifications (PTMs) and the presence of additional binding partners are almost certain to play important roles in regulating SMN oligomerization. In addition to Gemin2 and the Sm proteins, which bind directly to the Gem2 and Tudor domains of SMN, respectively, two other Gemin subcomplexes associate with the YG box. The Gemin3-4-5 and Gemin6-7-8 subunits are mainly tethered to SMN via Gemin3 and Gemin8, respectively [27]. Binding of these additional subunits to the oligomeric core of SMN·G2 will doubtless have an effect, but it is hard to predict its direction. Should these PTMs and binding interactions work to increase the local concentration of SMN, they would be predicted to drive demixing of the complex into phase-separated membraneless organelles [65] such as nuclear Cajal and Gemini bodies, or cytoplasmic stress granules and U bodies. Such a finding would also suggest that SMN might carry out different functions in different cellular locales and that these functions might depend on its oligomerization status. Thus, an important goal for the future is to elucidate the nature of the interactions that give rise to the underlying oligomeric heterogeneity of SMN subcomplexes *in vivo*. The work shown here not only demonstrates that formation of higher-order SMN complexes is required for metazoan viability, but it also provides key mechanistic insight into their assembly.

## Methods

### Protein Expression Purification and Reconstitution of SMN·Gemin2

Human and nematode SMN-G2 complexes were purified as previously described [33, 34]. SMN-Gemin2 complex was produced by co-expression of SMN (residues 1-294) and Gemin2 (12-280) fused to a C-terminal Mxe intein (NEB) containing a hexahistidine tag. Both coding regions were cloned into pETDuet (Novagen) and soluble SMN-Gemin2 complex was obtained following induction with 500 mM IPTG in BL21(DE3) cells for 16 h at 18°C. The complex was purified by Ni-NTA Superflow (Qiagen) and chitin bead (NEB) chromatography at 4°C, following the vendor’s protocols. SMN-Gemin2 complex was further purified on a Superdex 200 16/60 column at 20°C using 20 mM Tris·HCl, pH 7.5, 400 mM NaCl, 5 mM DTT, followed by spin column concentration (Millipore). SMNΔ5-Gemin2 complexes were produced using a similar protocol, except that SMNΔ5 was expressed from pCDFDuet and Gemin2-Mxe-His6 was produced from pETDuet together in BL21(DE3) and the final sizing column was performed using a Superdex 200 10/30 column at 4°C. Yeast SMN-G2 complexes were also purified as previously described [34].

The *Drosophila* SMN·Gemin2 complex was produced by separately expressing and purifying the two proteins and then reconstituting the complex *in vitro*. dmGemin2 was fused to a C-terminal Mxe intein (NEB) containing a hexahistidine tag, expressed in BL21(DE3) for 16 h at 18°C, and purified using Ni-NTA Superflow (Qiagen) and chitin bead (NEB) chromatography at 4°C following the vendor’s protocols. Purified protein was further purified on a Superdex 200 16/60 column at 20°C using 20 mM Na/KPO4 pH 7.0, 300 mM NaCl, and 10 mM β-ME. Similarly, dmSMN was fused with an N-terminal His7-Flag-Sumo (HFS) tag and cloned into a pCDFDuet expression vector and expressed in BL21(DE3) for two hours at 37°C following induction with 500 mM IPTG. Bacterial pellets were lysed in 100 mM NaKPO4, 10 mM Tris, 6M guanidine·HCL, 10 mM Imidazole, and 10 mM β-ME (final pH 8.0). After binding to Ni-NTA Superflow (Qiagen) resin, the column was subsequently washed with 15 c.v. of 100 mM Na/KPO4, 150 mM NaCl, 8M urea, 10 mM Imidazole, and 10 mM β-ME (final pH 7.7), followed by 20 c.v of 50 mM Na/KPO4 pH 7.0, 500 mM NaCl, 20 mM Imidazole, and 10 mM β-ME. Protein was eluted in 50 mM Na/KPO4 pH 7.0, 400 mM NaCl, 300 mM Imidazole, and 10 mM β-ME.

A two molar excess of purified dmGemin2 was added and the N-terminal HFS tag was liberated from SMN by overnight cleavage with SUMO protease Ulp1 (Life Sensors) at 4°C. After passage through a second Ni-NTA column to capture fusion protein, the reconstituted dmSMNG2 complex was purified on a on a Superdex 200 16/60 column at 20°C using 20 mM Na/KPO4 pH 7.0, 300 mM NaCl, and 1 mM DTT, followed by spin column concentration (Millipore).

### Size-Exclusion Chromatography In-line with Multiangle Light Scattering (SEC-MALS)

Analyses were performed as previously described [66]. Experiments were performed with a Superdex 200 10/300 GL column (GE Healthcare) at 0.5 ml/min at room temperature in 50 mM HEPES 7.5, 250 mM NaCl, and 5 mM DTT. The column was calibrated using the following proteins (Bio-Rad): thyroglobulin (670 kDa, R_S_ = 85 Å), γ-globulin (158 kDa, R_S_ = 52.2 Å), ovalbumin (44 kDa, R_S_ = 30.5 Å), myoglobin (17 kDa, R_S_ = 20.8 Å), and Vitamin B12 (1,350 Daltons). Blue-Dextran (Sigma) was used to define the void volume of the column.

Absolute molecular weights of the proteins studied were determined using multi-angle light scattering coupled in-line with size-exclusion chromatography. Light scattering from the column eluant was recorded at 16 different angles using a DAWN-HELEOS MALS detector (Wyatt Technology Corp.) operating at 658 nm. The detectors at different angles were calibrated using the monomer peak of Fraction V bovine serum albumin (Sigma). Protein concentration of the eluant was determined using an in-line Optilab T-rEX Interferometric Refractometer (Wyatt Technology Corp.). The weight-averaged molecular weight of species within defined chromatographic peaks was calculated using the ASTRA software version 6.0 (Wyatt Technology Corp.), by construction of Debye plots (KC/_*R*θ versus. sin^2^[θ/2]) at one second data intervals. The weight-averaged molecular weight was then calculated at each point of the chromatographic trace from the Debye plot intercept and an overall average molecular weight was calculated by averaging across the peak.

### Sedimentation Velocity

*S*edimentation velocity analytical ultracentrifugation (SV-AUC) experiments were performed at 20°C with an XL-A analytical ultracentrifuge (Beckman) and a TiAn60 rotor with two-channel charcoal-filled epon centerpieces and quartz windows. Experiments were performed in 20 mM HEPES-NaOH (pH 7.5), 300 mM NaCl, and 0.1 mM TCEP at concentrations of 0.16–1.0 mg/mL. Complete sedimentation velocity profiles were collected every 30 s for 200 boundaries at 40,000 rpm. Data were fit using the c(s) distribution model of the Lamm equation as implemented in the program SEDFIT [67]. After optimizing meniscus position and fitting limits, the sedimentation coefficient(s) and best-fit frictional ratio (*f*/*f*_0_) was determined by iterative least squares analysis. Sedimentation coefficients were corrected to *s*2o,w based on the calculated solvent density (ρ) and viscosity (η) derived from chemical composition by the program SEDNTERP [68].

### Mixed Oligomer Experiments

SMNΔ5oGemin2 was expressed and purified as described above, with an additional Mono Q ion exchange purification step. HFS-SMNΔ5 ·Gemin2 (wild-type, patient mutation variants, and a truncated SMNΔ5^14-156^ negative control) was expressed using a pColADuet-derived vector and purified in the same way. Using both expression vectors, the two complexes were also co-expressed and purified using the same Ni-NTA, Chitin Binding Resin. The final material was spin-concentrated in a final buffer of 20 mM HEPES pH 7.5, 400 mM NaCl, and 10 mM DTT. The recovered proteins were analyzed by 12.5% SDS-polyacrylamide gel electrophoresis and visualized using acidic Coomassie blue.

For mixing experiments, material was further purified using Mono Q ion exchange purification steps. Protein mixtures were nutated at 10 μM concentrations for one hour at 25°C or 4°C before binding to 0.5 mLs of Ni-NTA resin at room temperature. Resin was washed twice with ten column volumes of wash buffer (20 mM Na/KPO4 pH 7.4, 400 mM NaCl, 20 mM imidazole) before elution with 200 mM imidazole and subsequent SDS-PAGE analysis as described.

### Yeast complementation analysis

A haploid *S.pombe* strain (spGV40) with chromosomal *smn1+* replaced by a kanMX6 marker and episomal *smn1+* provided on a *ura4+* plasmid (pGV2887) was constructed using standard methods [69] from strain ATCC 96116 *(h+ his3-D1 leu1-32 ura4-D18 ade6-M210)* pGV2887 was constructed by insertion of a genomic pcr fragment extending ~500 bp upstream and downstream of the Smn1p coding sequence into the BamHI/SalI sites of pUR19 [70] using primers 5’-gtcagt-ggatccttactgagaagtcctcgctaaacc-3’ and 5’-gtcagtgtcgacatcaccaccgtggagacgaac-3’. Wild-type and mutant Smn1p coding sequences were cloned into pREP3X [71] and transformed into spGV40 with selection on Edinburgh minimal medium (Bio 101) supplemented with histidine, adenine, lysine, and thiamine but not leucine (EMMT-Leu). Six transformants were patched onto EMMT-Leu plates and replica plated to dilution on successive EMMT-Leu plates followed by FOA selection plates containing 0.5% yeast extract, 80 μg/ml adenine sulfate, 3% dextrose, 0.5 mg/ml 5-fluoroorotic acid (Research Products International), adjusted to pH 4 with acetic acid. Growth on FOA medium requires loss of the ura4+ marker, indicating that the pREP3X plasmid can support growth in the absence of *smn1+* from pGV2887. Experiments were repeated three or more times with consistent results for all constructs but the Y137H and Y137R mutants, which often showed more growth on FOA than shown in Fig 3B, but always much less than wild-type Smnp. Basal expression in the presence of thiamine was sufficient for complementation of the *smn1+* deletion.

### Drosophila husbandry, transgenesis and viability

Balanced transgenic fly lines (overall genotype: *Smn^X7^, Flag-Smn^TG^/TM6B-GFP)* were generated as previously described [29], where “TG” represents a given transgene. Briefly, the lines were generated using ΦC31 integration at an insertion site located in chromosome band position 86F8 and these lines were introgressed into an *Smn^X7^* null mutant background. The *Smn* transgenic construct is a ~3kb fragment containing the entire *Smn* coding region, expression of which is driven by the native *Smn* promoter. The transgene also contains an N-terminal 3X-FLAG tag. The *Smn^X7^* and *Smn^D^* alleles are previously described null alleles [54, 72], and both stable stocks are GFP-balanced. To generate singlecopy transgenic mutants *(Smn^X7^,Smn^TG^/Smn^X7^)*, virgin *Smn^X7^*/TM6B-GFP females were crossed to *Smn^X7^,Smn^TG^*/TM6B-GFP males. Crosses were performed on molasses-based agar plates with yeast paste, and then GFP-negative larvae were sorted into vials containing standard molasses fly food at the 2^nd^-instar stage to prevent competition from heterozygous siblings.

To measure viability, 25-50 GFP-negative progeny at the late 2^nd^ to early 3^rd^-instar stages were sorted into vials containing standard molasses fly food. After sufficient time had passed, pupal cases were counted and marked, and any adults were counted and removed from the vial. Any new pupal cases or adults were recorded every two days. The % viability was calculated at both the pupal and adult stages. Pupal viability (% pupation) was calculated by dividing the number of pupal cases by the initial number of larvae and multiplying by 100 (# pupae/# initial larvae*100). Adult viability (% eclosion) was calculated similarly but using the number of adults as the numerator (# adults/# initial larvae*100).

To assess larval motor function, crosses were maintained and progeny were raised at 25°C. Once the larvae reached the wandering 3^rd^-instar, 1-5 animals were placed onto the locomotion stage (a large molasses plate) at room temperature. The stage was then placed into a recording chamber to control light and reflections on the stage. Once all larvae were actively crawling, movement was recorded for at least 62 seconds on an iPhone6 at minimum zoom. Two recordings were taken for each set of larvae. At least 30 larvae were recorded for each experimental group. Locomotion videos were transferred to a PC and converted to raw video .avi files using the ffmpeg program. Videos were then opened in Fiji/ImageJ (https://imagej.net/Fiji), trimmed to about 60sec of video, and converted into a series of binary images. The wrMTrck plugin for ImageJ (http://www.phage.dk/plugins/wrmtrck.html) was used to analyze the video and determine larval size, average speed of movement, and average speed normalized to larval size (body lengths per second or BLPS) [73]. Each larva was treated as an individual when calculating average and standard error.

### Structural Modeling

A structural model of a dmSMN YG-box dimer was generated by replacement of side chains in the spSMN YG-box (PDB entry 4RG5) with those that differ in the dmSMN sequence. A hsSMN dimer model was extended to include additional N- and C-terminal residues using the same superposition. A model of an hsSMN tetramer was generated by superposition of one helix from each of two YG-box dimers with helices forming a right-handed cross in the *E. coli* glycerol facilitator structure (PDB 1FX8). Residues 269-277 of one hsSMN helix were superposed onto residues 92-100 of 1FX8 and the same residues of a second hsSMN helix were superposed onto residues 14-22 of 1FX8. The resulting interface is tightly packed and required adjustment of His273, Tyr277, and Met278 side chains. Further adjustment of the inter-helical distance and angle would be required to fully relieve the remaining steric contacts.

## Acknowledgements

Work in the Matera laboratory was supported by NIH grant R35-GM136435 (to A.G.M). K.G. acknowledges support from the Johnson Research Foundation. Additional support comes from the NIH project ALS-ENABLE (P30-GM124169) and a High-End Instrumentation Grant S10-OD018483. The SE-AUC, SV-AUC, and SEC-MALS experiments were performed at the Johnson Foundation Biophysical and Structural Biology Core Facility (University of Pennsylvania, Philadelphia PA). SAXS analyses were conducted at the Advanced Light Source (ALS), a national user facility operated by Lawrence Berkeley National Laboratory on behalf of the Department of Energy, Office of Basic Energy Sciences, through the Integrated Diffraction Analysis Technologies (IDAT) program, supported by DOE Office of Biological and Environmental Research. SEC-SAXS data were also obtained at 16-ID (LIX) at the National Synchrotron Light Source II, a U.S. Department of Energy (DOE) Office of Science User Facility operated for the DOE Office of Science by Brookhaven National Laboratory under Contract No. DE-SC0012704.

