## Supplemental Materials for "Assembly of higher-order SMN oligomers is essential for metazoan viability and requires an exposed structural motif present in the YG zipper dimer"

### Supplemental Methods

*Purification of MBP-dmSMN$^{189-220}$*. MBP-dmSMN$^{189-220}$ was expressed pETDuet at 37°C and purified on amylose resins (New England Biolabs) followed by Superdex-200 sizing (G.E. Healthcare). Proteins were stored in 20 mM Na/$KPO_4$ pH 7.0, 300 mM NaCl, 10 mM ß-ME, and 10% glycerol at -80$^o$C.

*Sedimentation equilibrium analysis (SE-AUC).* Analytical ultracentrifugation experiments were performed with an XL-A analytical ultracentrifuge (Beckman-Coulter) and a TiAn60 rotor with six channel charcoal-filled epon centerpieces and quartz windows. SE data were collected at 4$^o$C with detection at 280 nm for 1-3 sample concentrations in 20 mM Tris 7.4, 200 mM NaCl, 5 mM DTT. Analyses were carried out using global fits to data acquired at multiple speeds for each concentration with strict mass conservation using the program SEDPHAT (1). Error estimates for equilibrium constants were determined from a 1,000-iteration Monte Carlo simulation. The partial specific volume ($\bar{\upsilon}$), solvent density ($\rho$), and viscosity ($\eta$) were derived from chemical composition by SEDNTERP (2).

*Small-Angle X-ray Scattering at NSLS Beamline X21.* Sample scattering profiles from beam line X21 at the National Synchrotron Light Source (NSLS, Upton, NY, USA) were collected with a MAR 165 CCD detector (Mar USA, Inc., Evanston, IL). Two-dimensional images were integrated using software developed at the beam line into one-dimensional intensity profiles as a function of q. Measurements were taken at 20°C with a sample-to-detector distance of 835 mm and an X-ray wavelength of 10 kEV; scattering profiles covered a q range from 0.009 to 0.45 $Å^{-1}$. The sample holder was a 1-mm quartz capillary (Hampton Research, Aliso Viejo, CA) that was sealed across the evacuated beam path. Both ends of the capillary were open to allow the sample to flow continuously through to minimize radiation damage to the sample. Each measurement required 30 µl of sample for 30s and 60s exposure times. After each measurement, the capillary was washed repeatedly with buffer solution and purged with compressed nitrogen.

*Size-exclusion chromatography (SEC)-SAXS*. Data were collected at the SIBYLS beamline of the Advanced Light Source Light Source II (Berkeley, CA) (3). Data were collected at a wavelength of 1.0 Å in a three-camera conformation, yielded accessible scattering angle where $0.006 < q < 3.0$ $Å^{-1}$, where q is the momentum transfer, defined as $q = 4\pi \sin(\theta)/\lambda$, where λ is the X-ray wavelength and 2θ is the scattering angle; data to $q<0.5$ $Å^{-1}$ were used in subsequent analyses. 100 µL of 2.5 mg/mL dmSMN•G2 or 10 mg/mL MBP-dmSMN$^{189\text{-}220}$ were injected and eluted isocratically from a Shodex 804 sizing column equilibrated in 20 mM $N/KPO_4$ pH 7.0, 300 mM NaCl, and 1 mM DTT, at room temperature. Eluent from the column flowed into a 1 mm capillary for subsequent X-ray exposures at 1-s intervals. Plots of intensity from the forward scatter closely correlated to in-line UV and refractive index (RI) measurements.

*SAXS Analysis***.** SVD-EFA analysis of the SEC-SAXS data sets were performed as previously described (4), as implemented in the program RAW (5). Buffer subtracted profiles were analyzed by singular value decomposition (SVD) and the ranges of overlapping peak data determined using evolving factor analysis (EFA)(6). The determined peak windows were used to identify the basis vectors for each component and the corresponding SAXS profiles were calculated. When fitting manually, the maximum diameter of the particle ($D_{max}$) was incrementally adjusted in GNOM (7) to maximize the goodness-of-fit parameter, to minimize the discrepancy between the fit and the experimental data, and to optimize the visual qualities of the distribution profile. The theoretical SAXS profiles for atomic models were created using the FoxS program (8). The models were rendered using the program PYMOL (9).

*Parallel Axis Theorem Analysis* (10-12). Knowing the individual radii of gyration ($R_{1g}$ and $R_{2g}$) of two objects and their overall $R_g$ as a complex, the distance r between the two bodies can be expressed as such:

$$R_g{}^2 = f_1 R_{1g}{}^2 + f_2 R_{2g}{}^2 + f_1 f_2 r^2$$

where

$$f_i = \frac{\int \rho_i dV_i}{\int \rho_i dV_1 + \int \rho_i dV_2} \; (i=1,2)$$

These are the relative scattering components of the two particles, with respective scattering-length densities $\rho_i$ and volumes $V_i$.

**Table S1. Oligomeric Properties of SMN•G2 Chimeras.**

| SMN•G2 Construct | SEC-MALS $M_w$ (20°C)[B] | Peak Concentration (µM)[C] | Oligomer State |
|---|---|---|---|
| *H.sapiens* | ~247 kD (200-600) | | Tetramer-Octamer[A] |
| *H.sapiens* Δ5 | ~231 kD (188-454) | 0.7 | Tetramer-Octamer |
| SMNΔ7 | ~66 kD (58-74) | | Monomer[A] |
| *S.pombe* | ~150 kD (110-180) | | Dimer-Tetramer[A] |
| *C.elegans* | ~326 kD (250-450) | 0.2 | Tetramer-Octamer |
| *D.melanogaster* | ~345 kD (202-380) | 0.4 | Tetramer-Octamer |
| $Hs^{1-252}$-$Sp^{117-152}$ | ~264 kD (223-409) | 1.0 | Tetramer-Octamer |
| $Hs^{1-275}$-$Sp^{140-152}$ | ~128 kD (88-227) | 2.3 | Dimer |
| $Hs^{1-275}$-$Sp^{140-152\Delta Cys}$ | ~125 kD (117-132) | 1.5 | Dimer |
| $Hs^{1-279}$-$Sp^{144-152}$ | ~285 kD (119-640) | 0.1 | Tetramer-Octamer |
| $Hs^{1-275}$-GCN4(IL) | ~113 kD (80-113) | 0.3 | Dimer |
| $Hs^{1-251}$-GCN4(LI) | ~190 kD (190-400) | 0.0 | Tetramer-Octamer |
| $Sp^{1-117}$-GCN4(IL) | ~85 kD (79-85) | 2.1 | Dimer |
| $Sp^{1-117}$-GCN4(IL)-$spSMN^{145-152}$ | *n.d.* | *n.d.* | Dimer[A] |
| $Sp^{1-117}$-GCN4(LI) | ~125 kD (90-128) | 1.1 | Dimer-Trimer |
| $Sp^{1-117}$-GCN4(II) | ~113 kD (73-120) | 0.7 | Dimer-Trimer |
| $Sp^{1-141}$-$Ce^{187-197}$ | ~160 kD (120-170) | 0.5 | Dimer-Tetramer |
| $Ce^{2-184}$-$Sp^{140-152}$ | ~137 kD (137 – 610) | 0.9 | Dimer |

[A]See ref. (13)
[B]Shown is the MALS molecular weight determined at UV peak. In parentheses, the range of molecular masses observed from UV peak half-height to half-height is shown.
[C]As determined by refractive index.

**Table S2. Biophysical Properties of *sp*SMN•Gemin2 Mutants (related to Figure 3)**

M = Monomer, D = Dimer, T = Tetramer, O = Octamer, SS = Single Species

| | Sedimentation Equilibrium Analytical Ultracentrifugation[A] | | | | | SEC-MALS[C] |
|---|---|---|---|---|---|---|
| Construct | Speeds (krpm) | Conc. (μM) | Model of Association | $K_d$ (μM) | $X^2$ | Mass Profile |
| Wild-type[B] | 8,10,12,14 | 2.2, 3.3, 4.3 | 2-4 | 1.0 ± 0.9 | | D-T |
| Wild-type | 8,10,12,14 | 2.3 | 2-4 | 2.8 ± 0.4 | 0.9 | D-T |
| S130Q | 12,14,16 | 5.6, 3.6 | 2-4 | 4.0 ± 0.25 | 1.4 | D>T |
| A134Q | 12,14,16 | 9.2, 7.9 | 2-4 | 14.3 ± 0.09 | 0.6 | D |
| Y136C | 8,10,12,14 | 6.5 | SS 1:1 | *n.a.* | 0.9 | M |
| Y137R | 8,10,12,14 | 2.2 | 2-4 | 274 ± 32 | 1.0 | D |
| T138I | *n.d.* | *n.d.* | *n.d.* | *n.d.* | *n.d.* | M+ |
| G139S | 8,10,12,14 | 1.2 | 1-2 | 4.5 ± 0.75 | 0.4 | M-D |
| L140Y,A141Y | *n.d.* | *n.d.* | *n.d.* | *n.d.* | *n.d.* | T-O |
| A141Q | 12,14,16 | 6.3, 4.6 | 2-4 | 0.55 ± 0.07 | 1.7 | D-T |
| E142R | *n.d.* | *n.d.* | *n.d.* | *n.d.* | *n.d.* | D-T |
| G143V | *n.d.* | *n.d.* | *n.d.* | *n.d.* | *n.d.* | D-T |
| A145Q | 12,14,16 | 3.5 | 2-4 | 1.1 ± 0.3 | 0.9 | D-T |
| spSMN$^{1\text{-}141}$-ceSMN$^{187\text{-end}}$•Gemin2 | *n.d.* | *n.d.* | *n.d.* | *n.d.* | *n.d.* | D-T |
| spSMN(GCN4IL)•Gemin2 | 8,10,12,14 | 1.4 | SS 2:2 | *n.a.* | 2.1 | D |

*n.d.* = not determined

*n.a.* = not applicable

A. Analyses were performed in 20 mM Tris 7.4, 200 mM NaCl, 5 mM DTT and data analyzed using the program SEDPHAT (1).

B. in 20 mM Na/$KPO_4$ pH 7.0, 150 mM NaCl, 1 mM DTT (13).

C. Analyses were performed in 20 mM Tris 7.4, 200 mM NaCl, 5 mM DTT with a Superdex 200 10/300 column at room temperature.

**Table S3. Parameters derived from Size-Exclusion Chromatography In-line with Small-Angle X-ray Scattering (SEC-SAXS)**

| Sample[1] | Guinier | | GNOM | | [2]$P_x$ | Mass (kD) | | Oligomer |
|---|---|---|---|---|---|---|---|---|
| | $qR_g$ | $R_g$ (Å) | $R_g$ (Å) | $D_{max}$ (Å) | | [3]$Q_r$ | [4]Porod | |
| dmSMN•Gemin2 | 0.71 – 1.85 | 82.0 ± 0.8 | 83.7 ± 0.9 | 271 | 2.7 | 472 | 390 | Octamer |
| MBP-dmSMN$^{189-220}$ | 0.54 – 1.14 | 56.1 ± 0.7 | 53.0 ± 0.2 | 179 | 4.0 | 345 | 409 | Octamer |
| | 0.56 – 1.21 | 38.0 ± 0.4 | 41.2 ± 0.1 | 113 | 4.0 | 239 | 227 | Tetramer |

[1]In 20 mM Na/$KPO_4$ pH 7.0, 300 mM NaCl, 1 mM DTT
[2]Porod exponent. Values near ~4 indicate compactness, whereas lower values between <2-3 indicate significant lack of compactness and increased volumes (14). These values were determined using the program ScÅtter (https://bl1231.als.lbl.gov/scatter/).
[3]Mass determinations (MM) using the $Q_r$ invariant (15) were determined using the program RAW(5).
[4]Porod Volume ($V_p$). This figure can be used to estimate the mass of compact proteins, where $V_p/1.6 \sim$ MM.

**Table S4. Parameters derived from Small-Angle X-ray Scattering (SAXS) for spSMN(GCN4IL)•Gemin2**

| Conc.[1] | Guinier | | GNOM | | [2]$P_x$ | Mass (kD) | | Oligomer |
|---|---|---|---|---|---|---|---|---|
| (mg/mL) | $qR_g$ | $R_g$ (Å) | $R_g$ (Å) | $D_{max}$ (Å) | | [3]$Q_r$ | [4]Porod | |
| 12 | 0.52 – 1.10 | 65.1 ± 0.4 | 67.5 ± 0.3 | 233 | 3.5 | 113 | 126 | Dimer |
| 6 | 0.52 – 1.10 | 65.3 ± 0.7 | 68.7 ± 0.3 | 229 | 3.7 | 115 | 133 | Dimer |
| 4 | 0.57 – 1.15 | 63.9 ± 0.8 | 66.8 ± 0.3 | 219 | 3.6 | 110 | 127 | Dimer |
| 3 | 0.57 – 1.15 | 63.8 ± 1.1 | 65.7 ± 0.4 | 199 | 3.9 | 106 | 121 | Dimer |

[1]In 20 mM Na/$KPO_4$ pH 7.0, 150 mM NaCl, 1 mM DTT
[2]Porod exponent. Values near ~4 indicate compactness, whereas lower values between <2-3 indicate significant lack of compactness and increased volumes (14). These values were determined using the program ScÅtter (https://bl1231.als.lbl.gov/scatter/).
[3]Mass determinations (MM) using the $Q_r$ invariant (15) were determined using the program RAW(5).
[4]Porod Volume ($V_p$). This figure can be used to estimate the mass of compact proteins, where $V_p/1.6 \sim$ MM.

YG Box: Extended Phylogenetic Comparison

```
                         ————————— YG Box —————————
Human         DDADALGSMLISWYMSGYHTGYYMGFRQNQKEGRCSHSLN
Dog           DDADALGSMLISWYMSGYHTGYYMGFKQNQKEGRCSHFN
Pig           DDADALGSMLISWYMSGYHTGYYMGFRQNQKEGRCSHFN
Mouse         DDTDALGSMLISWYMSGYHTGYYMGFRQNKKEGKCSHTN
Chicken       EDDEALGSMLIAWYMSGYHTGYYLGLKQSRMEAALEREAYLK
Frog          EDDEALGSMLISWYMSGYHTGYYLGLKQGRMESSIGKPPHQK
Gator         EDDEALGSMLIAWYMSGYHTGYYLGLKQSRMEAALDRHPDPK
Snake         DDDEALGSMLIAWYMSGYHTGYYLGLKQGRMEATLERHAHSK
Gekko         EDDEALGSMLIAWYMSGYHTGYYLGLKQSRMEATSGRDAHSK
Killifish     VDDEALGSVLISWYMSGYHTGYYLGLKQGRKEANKWTKLHHK
Zebrafish     EDDEALGSMLISWYMSGYHTGYYMGLRQGRKEAAASKKSHRK
Fugu          VDDEALGSMLISWYMSGYHTGFYLGLKEGRKKASNWTKPHHR
Arowana       SDVAELSSMLLSWYLCGYHTGYYMALQQTNSSHEKTKKKYK[10aa]
Catfish       EDDEALGSMLISWYMSGYHTGYYLGLKQGRKEAAAKKSHYK
Shark         EDDEALGSMLIAWYMSGYHTGYYLGLKQGRAEEALGKSSHRK
Coelacanth    DADSTLVCMLLAWYMSGYHTGYYMGLKHGQAKATGSSQKKHPKRK
Octopus       DNNEALCSMLMSWYMSGYHTGYYQGLKSKQN
Urchin        MDKEALHSMLMSWYMSGYHTGYYEGMKKSKTSSHSATSKPK[42aa]
Oyster        GDNEAMCSMLMAWYMSGYHTGYYQGLKQGRQQGGTSPHPDSFR
Mollusk       GDNEAMCSMLMAWYMSGYHTGYYQGLKQGRQQGGTSPHPDSFR
Anemone       EDNEALASMLMSWYLSGYYTGYYQGTQRQHISHNDRTSQTT[23aa]
Coral         HDNDALASMLMAWYLSGYHTGYFQAMQNFRHESSMGSNEAQ[19aa]
Hydra         GDEEALAGMLMSWYMSGYHTGYYQGMHFLNRDSESKKIKNN[97aa]
Ciona         LSKDALSNMFASWYMAGYQTGFHRGLASSCCKNNCKK
Crab          TDDEALASMLMSWYMSGYHTGYYQALRQMRSESCECQGHIDKKCCLHCSS
Daphnia       MDSDSLYSMLMSWYMAGYHTGYYQGVQQRNKRKANSSGSS
Psyllid       DTTESLSAVLMAWYMAGYHTGRYEASLGLNRNKTFGRPQSQQVEKCCDHK
Bug           DESDALSAMLMAWYMSGYHTGYYQGLTRSTPSGSGIREGKSNTSRLNN
WaterTick     KESDALSSMLMAWYMSGYHTGYYQALQQSQQVEKCCDHK
Mite          EGDEALAAMLISWYISGYHTGYYTAVRNQSKG
Aphid         TEREALTSMLMSYYMSGFHTGYYLGIKQKNSEN
Ant           TDADALSSMLMSWYLSGFHTGYYHGLKQAKNQQKRRNC
Spider        SNDPSLSAMLVSWYMAGYYTGLHQVCI
Wasp          NDADALSSMLMSWYISGFHTGYYHGLKQAQSNQHRRKT
Honeybee      NDAEALSSMLMSWYISGFHTGYYHGLKQAEKNQTKRKNC
Mosquito      VESENLSAMLMSWYMSGYYTGLYHGQRMSQQQQHTQQKRARQS
Silkmoth      SEQQAISSMLLSWYMSGYYTGLYQGMKRSKENNKNV
Housefly      EDSEHLSAMLMSWYMSGYYTGVYQGMQMAKSKTKK
D.pseudo      GEEQDLMSMLTAWYMSGYYTGYFQGKKAIPRQVEKKKTPKK
D.melano      GAEQDFVAMLTAWYMSGYYTGLYQGKKEASTTSGKKKTPKK
Nematode      NQKEAMNSMLMSWYMSGYHTGYYQGLADQKNVQN
HookWrm       DEAEALSSMLMAWYMSGYHTGYYQAIRDMNSNS
FlatWrm       NDENAVRNLLNSWYMCGYQTGYLTALKKGARECV
TapeWrm       NDENAVRNLLNSWYMCGYQTGYLTALKKGTRECV
WhipWrm       KVDNALNAMLISWYMAGYHAGFYQAGTHSWFVMAYGFSVCSG[36aa]
AcornWrm      GDEDALFSMLISWYMSGYHTGYYQGLKASRGESLQNTPQRNE[24aa]
Priapulus     IDNDVLSSMLISWYMSGYHTGYYQGLKTAKLEALKKRKDDVP[27aa]
Lingula       GDNEALCSMLMSWYMSGYHTGYYQGLKDAKKQQHGHQPHKDSSDVKR
Placozoa      INDELLSNLIMSWYMAGYRTGYYQAIRKAKDKSKKK
S.rosetta     AQEAALANMLMSWYQSGFYTGYYQALQQLQQQQQE
Dicty         QGDDELADLLLSWYYSGYYTGVYQERKRNSRLNQTPHPNHIN[69aa]
Neurosp       VQDEELKKLLMSWYYAGYYTGLYEGKQKALHEQAQQ
B.bassia      GRDDNLRKLLMSWYYAGYYTGLHEGQQQQTQAQQQPQ
S.pombe       TYDETYKKLIMSWYYAGYYTGLAEGLAKSEQRKD
P.italic      VQDESLKNLMMSWYYAGYYTGLHEGQQQTNSNQSS
Trypano       RLPADIRQLLVAYFNAGYEAGYYVGKRDGSSKVGGKRARGE
Cotton        SCETDLTVLLNAWYSAGFYTGKYLVEQSIAKRRQ
Corn          NLDSDLAALLNSWYAAGFYTCRYLMLQSTKNSRP
Legume        DSATDLTAVLNAWYSAGFHTGKYLAEQSIGNRRQI
Jute          TSETDLTVLLNAWYSAGFHTGKYLMEQSIAKKQHK
Consensus          LxSMLxSWYxSGYxTGYxxGL
```

**Figure S1A (related to Figure 1). Extended phylogenetic alignment of the SMN YG Box.** Phylogenetic analysis of SMN C-termini from a wide variety of eukaryotes. Conserved glycine residues are shaded in magenta, hydrophobic residues are in green, and polar residues in teal.

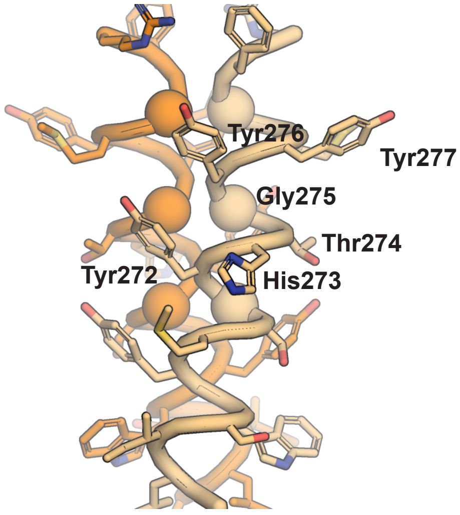


Human YG Box
PDB 4GLI

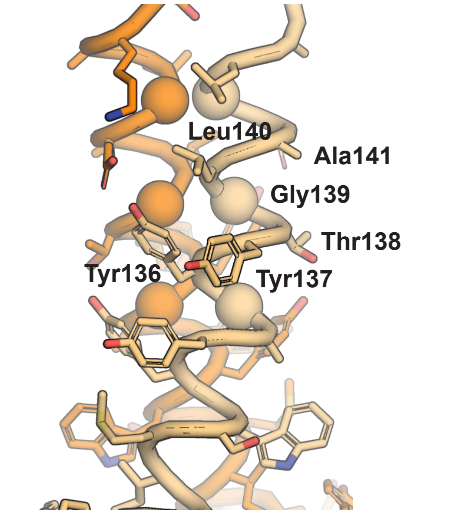


*S.Pombe* YG Box
PDB 4RG5

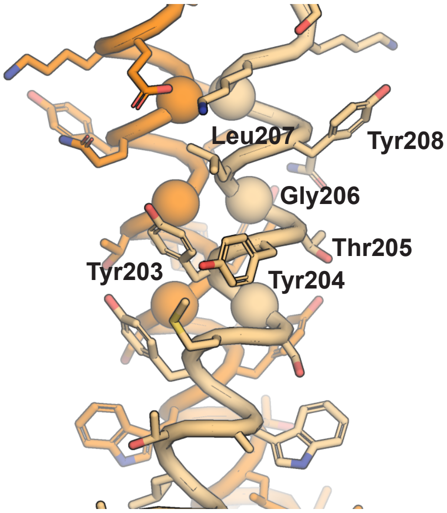


Model of *D. Melanogaster*
YG Box

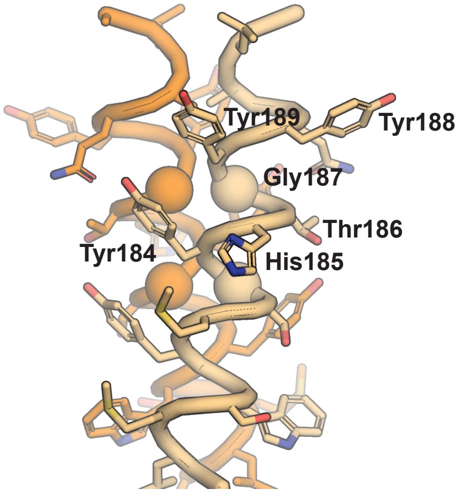


Model of *C. Elegans*
YG Box

**Figure S1B (related to Figure 1)**. Atomic views of the SMN YG-box dimer. Shown in the upper panels are the experimental atomic structures of the human (left, PDB 4GLI (16)) and worm (right, PDB 4RG5 (13)) YG boxes. In the lower panels are models of the fly (left) and worm (right) derived from the 4GLI structure. Shown as spheres in each panel are well-conserved glycine residues at $C_\alpha$. The residues corresponding to amino acids 272-277 in human are labelled in each panel. The figure was rendered using the program PYMOL (9).

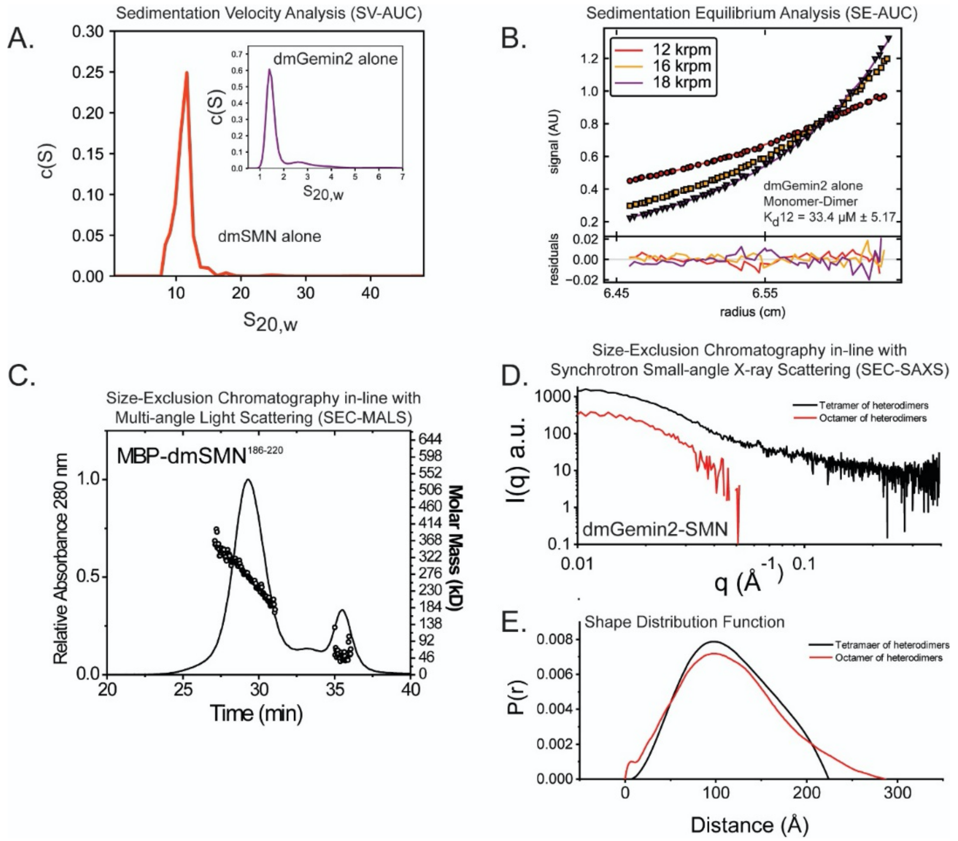


**Figure S2 (related to Figure 2) Biophysical characterizaton of *Drosophila* SMN and Gemin2.** ***A***. Sedimentation velocity analysis of dmSMN and dmGemin2 at 20°C. c(S) distributions derived from the fitting of the Lamm equation are shown for dmSMN (red) and dmGemin2 (blue, inset panel). This analysis of dmSMN shows evidence of a large ~11S species consistent with studies of the SMN•G2 complex from different species. The analysis of dmG2 alone shows evidence of a mostly monomeric species with some evidence of an oligomer. ***B***. Sedimentation equilibrium analysis of dmGemin2 at 4°C. Shown in the upper panels are radial absorbance data plotted with fitted models shown as solid lines for each of three speeds. Shown in the respective lower panels are the residuals for each fit. Data were best described with a monomer-dimer fit with a determined $K_d$ of 33.4 μM ± 5.2. C. SEC-MALS analysis of MBP-dmSMN$^{186\text{-}220}$. Shown as a black line is the absorbance profile of protein as a function of retention time in a Superdex-200 10/300 column at room temperature (left axis). Black circles denote molecular masses determined by in-line light scattering (right axis). The mass profiles determined span from dimer to octamer in the first peak and monomer in the second peak. D. SEC-SAXS analysis of dmSMN•G2. E. Resulting deconvoluted scattering profiles from SVD-EFA analysis of SEC-SAXS data collected for dmSMN•G2. Mass calculations using $Q_r$ and Porod volume relationship confirm the presence of tetrameric and octameric species. Model-independent properties derived from this analysis are shown in Supplemental Table 3. F. Shape Distribution Function analysis of tetramers (black) and octamers (red) of dmSMN•G2.

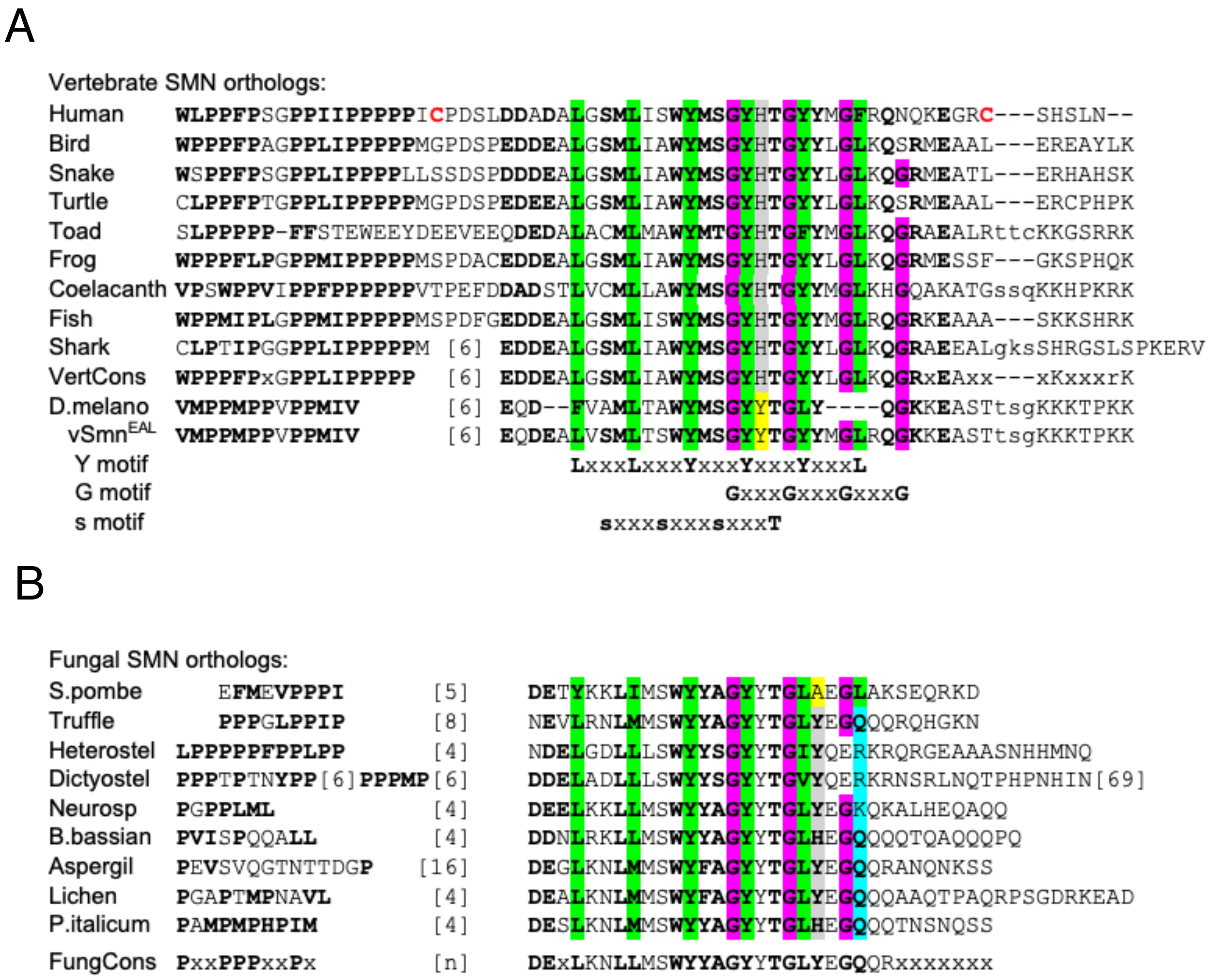


**Figure S3. Alignment of YG box sequences from vertebrate and fungal SMN orthologs.** ***A.*** Comparison of SMN YG boxes from nine different vertebrate clades showing the overall vertebrate consensus (VertCons). The three sequence motifs (Y, G and s) identified in Fig. 1A are shown below for comparison. Note the apparent conservation of a fourth GxxxG repeat (within context of QGRxE) among the vertebrates that provides motivation for insertion of MGLR sequence into *Drosophila* (D.melano) SMN to create vSmn and vSmn$^{EAL}$ (relates to Fig. 5A). A conserved histidine (His273 in human) is shaded in gray. ***B.*** Comparison of nine different fungal orthologs, revealing the overall lack of conservation of Ala141 (highlighted in yellow, relates to Fig. 6A). The fungal consensus sequence (FungCons) is presented below.

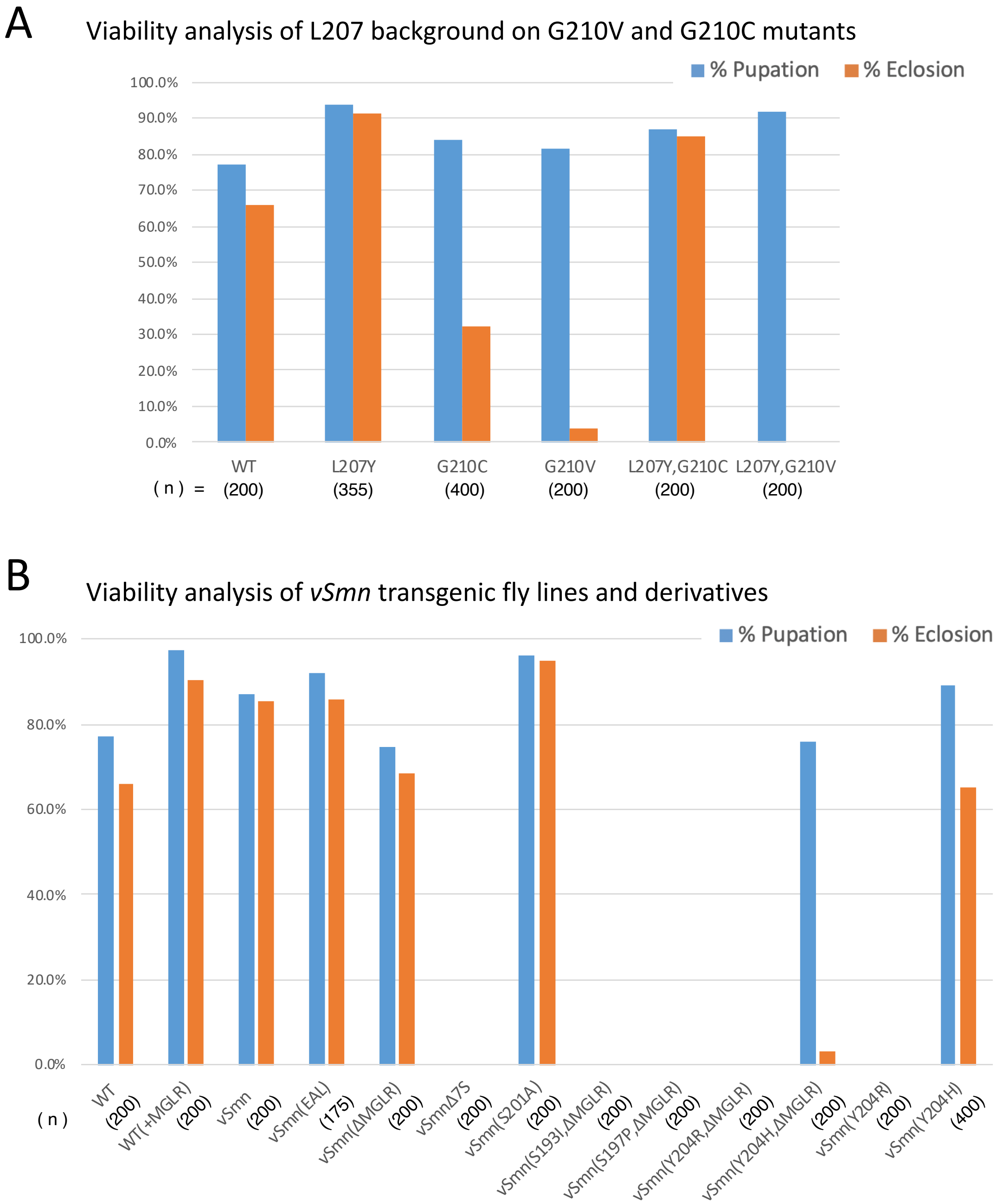


**Figure S4 (relates to Fig 5). Pupal and adult viability analysis of Flag-Smn transgenic lines.** ***A.*** Comparison of pupation and eclosion frequencies of wild-type (WT) and Smn missense mutations (relates to Fig. 5A). ***B***. Comparison of pupation and eclosion frequencies of vertebrate Flag-Smn (vSmn) fly lines as compared to WT controls (relates to Fig. 5A). For panels *A* and *B*, the total number of animals scored for each fly line is shown in parentheses below the genotype.

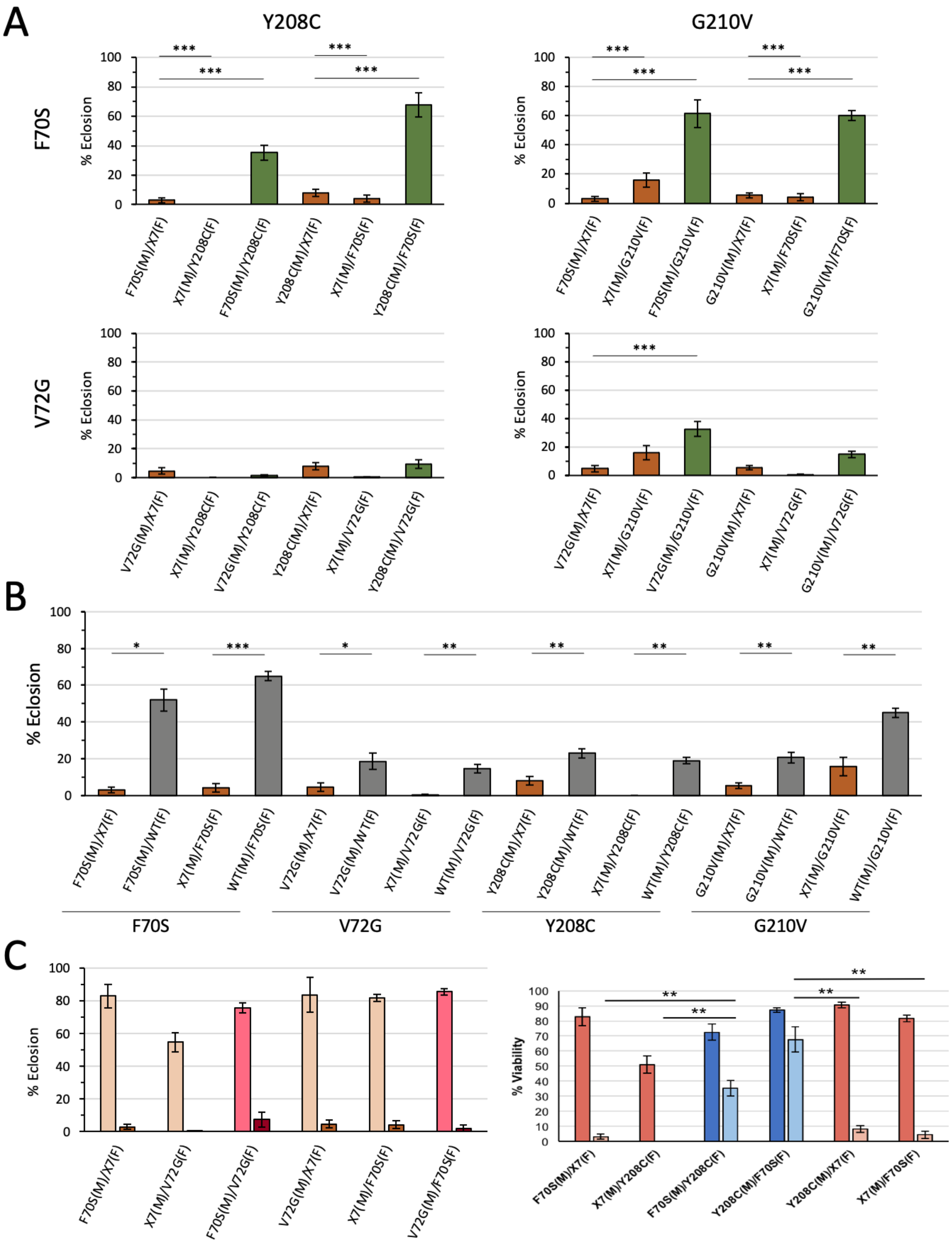


**Figure S5 (relates to Fig 6E). Intragenic complementation analysis.** ***A-C.*** Adult eclosion frequencies of various combinations of SMA-causing point mutations in either the tudor domain (F70S and V72G) or the YG box (G210V and Y208C). The genotype of each parental strain (male, M or female F) is shown in parenthesis. For each cross n>200 animals. Experiments carried out as previously described (see (17, 18)).

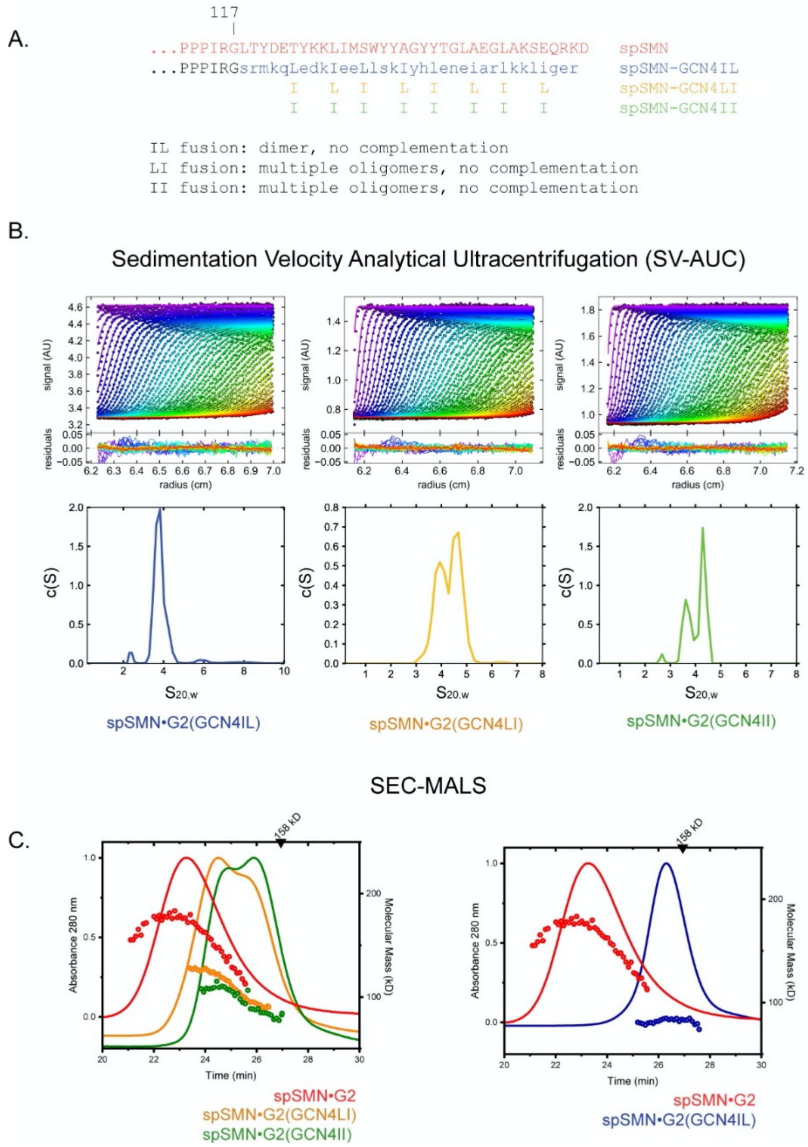


**Figure S6 (related to Figure 3). Analysis of spSMN-GCN4 chimeras.** ***A***. GCN4 variants fused to spSMN•G2. Shown are the GCN4 variants substituted for YG box regions spSMN•G2 constructs, along with their respective behaviors in yeast complementation assays. ***B***. Sedimentation velocity analysis. For each construct, representative absorbance data (colored circles) for sedimentation boundaries are shown in the upper panels as a function of radial position and time. Shown in solid lines are the fits to the Lamm equation, as performed in SEDFIT. In the respective lower panels, the residuals of these fits are shown. c(S) distributions derived from the fitting of the Lamm equation are shown for each. This analysis shows evidence of mostly dimers of GCN4(IL) constructs, while GCN4(LI) (yellow) and GCN4(II) (green) constructs show behavior consistent with the occurrence of trimers and tetramers. ***C***. SEC-MALS analysis. Shown for each construct as a black line is the absorbance profile of protein as a function of retention time in a Superdex-200 10/300 column at room temperature (left axis), each injected at 8 mg/mL. Black circles denote molecular masses determined by in-line light scattering (right axis). Relative to wild-type complex (red), GCN4(IL) fusions (blue) are dimeric, while the mass profiles for GCN4(LI)

(yellow) and GCN4(II) (green) constructs show evidence for higher order species approaching tetramer. All analyses here were performed in 20 mM Na/$KPO_4$ pH 7.0, 300 mM NaCl, and 2 mM DTT.

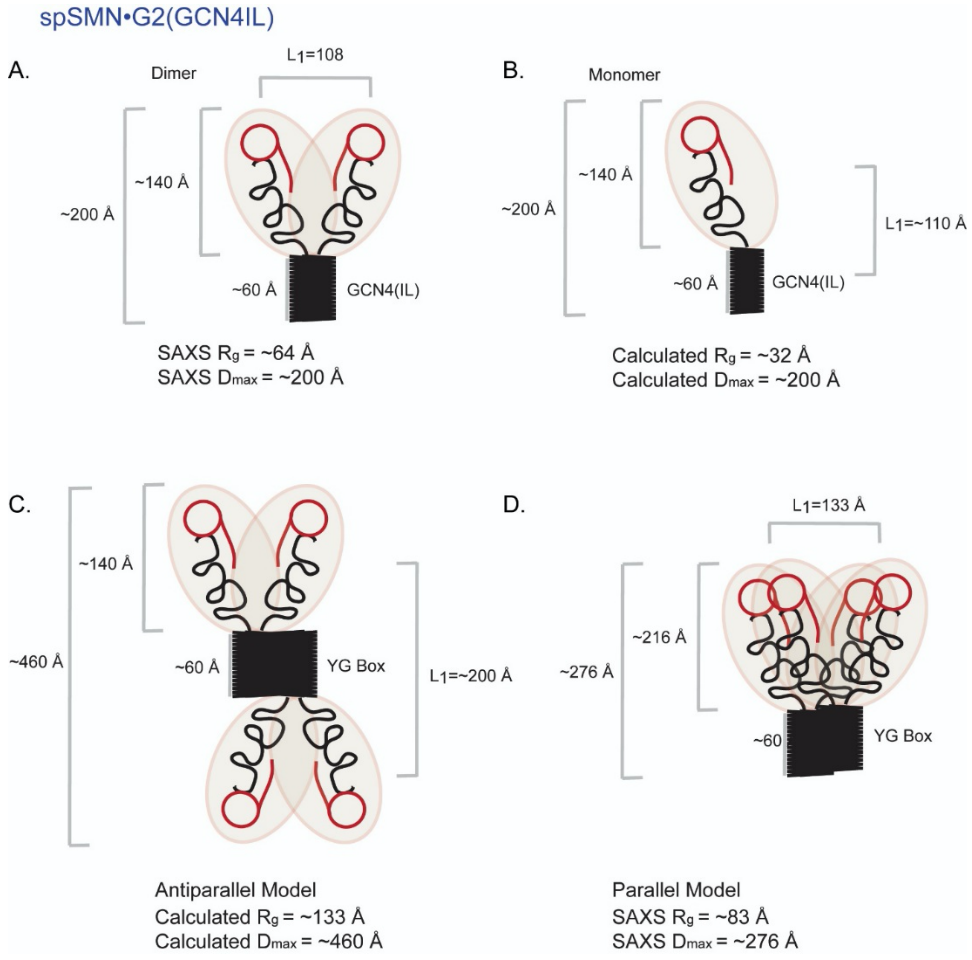


**Figure S7. Parallel Axis Theorem Analysis.** ***A-B***. Assuming that the maximum dimension of a spSMN-GCN4•G2 monomer and dimer are approximately the same, it is possible to approximate the $R_g$ of a monomer (~32 Å) using the relationship provided in Supplemental methods and the SAXS data presented in Table S4. The length of the YG box is ~60 Å as observed by X-ray crystallography. These figures, in turn, can be used to estimate the $L_1$ (center-to-center distance) for the spSMNGCN4•G2 dimer of ~108 Å. ***C-D***. spSMN(GCN4)•G2 dimer for the parameters of a dimer component in a tetramer, we can then consider two models for tetramerization. The calculated properties of such an antiparallel dimer in panel C are $R_g$ =133 Å and $D_{max}$= 460 Å, which is significantly larger than that experimentally determined for the spSMN•G2 complex (13) of $R_g$ =83 Å and $D_{max}$= 276 Å. Using this information, an $L_1$ of ~133 Å is calculated, more consistent with a parallel dimer.

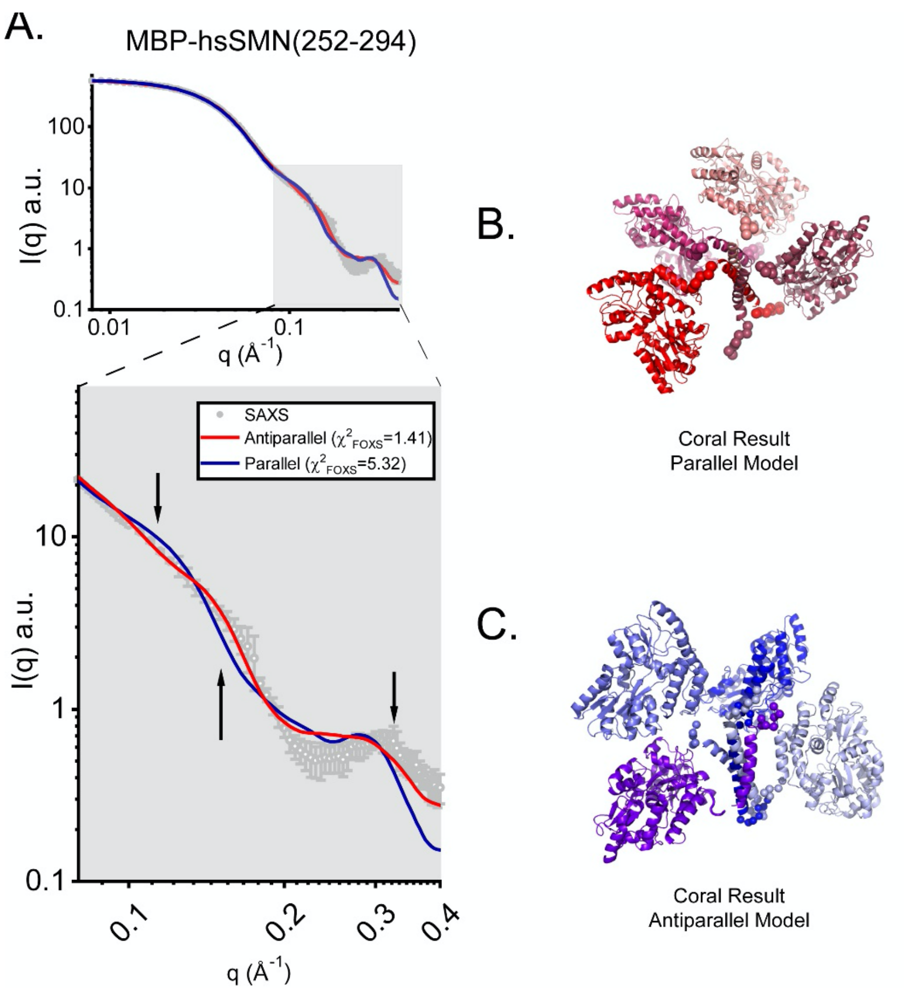


**Figure S8. CORAL analysis of MBP-hsYG$^{252\text{-}294}$.** In previous studies, MBP-hsYG$^{252\text{-}294}$ was extensively characterized using SAXS (16). Using these data, two models of the SMN tetramer were tested: a parallel dimer-of-dimers model (red) and an antiparallel mode (blue). In the current analysis, the YG components were fixed and the positions of the MBP domains and the flexible linker (residues 252-255, shown as beads) were refined in the program CORAL (19). Shown in panel A is the SAXS data against the calculated scattering for both models, as calculated by the program FoxS (8). In this analysis, the best agreement is observed for the parallel model ($\chi^2$=1.4) over the antiparallel model ($\chi^2$=5.3). Results were consistent across ten independent calculations in both cases. Representative results from CORAL analysis are shown in panels *B* (parallel model, red) and *C* (antiparallel model, blue).
